# CKII-Phosphorylated HPV-16 E7 Disrupts Planar Cell Polarity by Recruiting Vangl1 Phospho-E7 Hijacks Vangl1 Trafficking

**DOI:** 10.64898/2026.02.25.707969

**Authors:** Ifeoluwa D. Gbala, Om Basukala, Michael P. Myers, Rebecca Bertolio, Giannino Del Sal, Lawrence Banks

## Abstract

High-risk human papillomaviruses (HPVs) depend on continuous expression of the E6 and E7 oncoproteins to sustain malignant transformation. The high-risk E7 oncoprotein is multifunctional, and its phosphorylation by Casein Kinase II (CKII) amplifies its oncogenic potential. Here, we identify Vangl1, a core planar cell polarity (PCP) scaffold protein, as a novel, phosphorylation-dependent interactor of HPV-16 E7. CKII-mediated phosphorylation is required for Vangl1 recruitment, and this interaction is largely selective for HPV-16 E7. In cervical cancer cells, E7 expression profoundly disrupts Vangl1 homeostasis, producing a biphasic proteostatic imbalance in which phosphorylated Vangl1 is aberrantly retained. This retention parallels E7 stability, revealing a reciprocal oncogenic stabilization. E7 also impairs Vangl1 trafficking and localization by co-opting the clathrin adaptor subunit AP1M1. Functionally, Vangl1 depletion mirrors E6/E7 loss in CaSki spheroids by disrupting spheroid architecture, reducing invasiveness, and increasing chemosensitivity. Taken together, these findings position Vangl1 as a central effector in E7-mediated cervical transformation and invasive progression.

**Teaser:** E7’s control of Vangl1 reveals a polarity-remodeling mechanism that drives cervical cancer invasion and progression.

## Introduction

Cervical cancer remains a major global health challenge, disproportionately affecting women in low- and middle-income countries where access to screening, vaccination, and advanced treatment is limited(*1*, *2*). Despite the availability of effective prophylactic vaccines, persistent infection with high-risk human papillomaviruses (HPVs), particularly HPV-16 and HPV-18, continues to drive the majority of cervical cancer cases worldwide (*3*). These infections initiate a multistep progression from precancerous lesions to invasive carcinoma(*4–6*), a process sustained by the viral oncoproteins E6 and E7, which remain expressed throughout tumour development and are essential for malignant maintenance(*7–11*).

The HPV-16 E7 oncoprotein is a central driver of oncogenesis. Beyond its canonical inactivation of the retinoblastoma protein (pRB) and deregulation of E2F-dependent cell-cycle entry (*12–14*), E7 undergoes post-translational modifications that expand its oncogenic repertoire. Phosphorylation of conserved serine residues by casein kinase II (CKII) enhances E7 stability and reshapes its interactome (*15*, *16*), yet how phospho-E7 engages additional host pathways to promote malignancy remains incompletely understood. This gap is particularly important because HPV-positive tumours often display distinctive patterns of aggressive invasion and metastatic spread, suggesting that viral oncoproteins may influence cellular polarity and tissue organization(*17–20*).

Emerging evidence points to the non-canonical Wnt/planar cell polarity (PCP) pathway as a critical regulator of epithelial architecture, directional migration, and metastatic behaviour (*21–24*). Dysregulated PCP signalling disrupts polarity, activates Rho GTPases and JNK, and promotes epithelial-to-mesenchymal transition, collectively enabling invasive phenotypes across multiple cancers (*24–27*). Core PCP components, including Vangl1 and Vangl2, have recently been implicated in tumour progression, yet their involvement in HPV-driven malignancy remains largely unexplored. Prior studies show that HPV E6/E7 can activate PCP/JNK signalling (*25*, *26*), but the upstream viral–host interactions that initiate this activation are not known.

This unresolved link between HPV oncoproteins and PCP machinery raises a fundamental question of how HPV-16 E7 directly engages PCP components to reprogramme epithelial behavior and promote invasion. Here, we identify Vangl1 as a novel interactor of phosphorylated HPV-16 E7, and demonstrate that this interaction drives Vangl1 dysregulation in cervical cancer cells. Thus, Vangl1 is a compelling candidate for this missing link, given its central role in establishing planar polarity and its oncogenic functions in migration, metastasis, and therapy resistance (*28*). We show that phospho-E7–mediated perturbation of Vangl1 enhances invasive and collective metastatic phenotypes and alters chemotherapeutic responses in three-dimensional culture systems. These findings uncover a new mechanism by which HPV-16 co-opts PCP signalling to promote malignant progression and highlight Vangl1 as a potential therapeutic target in HPV-associated cancers.

## Results

### Phosphorylated HPV-16 E7 recruits Vangl1

Phosphorylation of high-risk HPV E7 oncoprotein by Casein Kinase II (CKII) at serine residues 31 and 32/33 is essential for maintaining the transformed phenotype in cervical cancer–derived cell lines (*49–51*). Given the importance of this modification, we sought to identify phospho-specific binding partners of HPV-16 E7 and to determine the functional relevance of such interactions. To this end, we synthesized biotin-tagged peptides corresponding to the CKII site within the CR2 region of HPV-16 E7 (Figure 1a), in both phosphorylated and non-phosphorylated forms (Figure 1b). The peptides were then used as bait in streptavidin pulldown assays with HaCaT keratinocyte lysates; the bound proteins were analysed by mass spectrometry.

**Figure 1:**
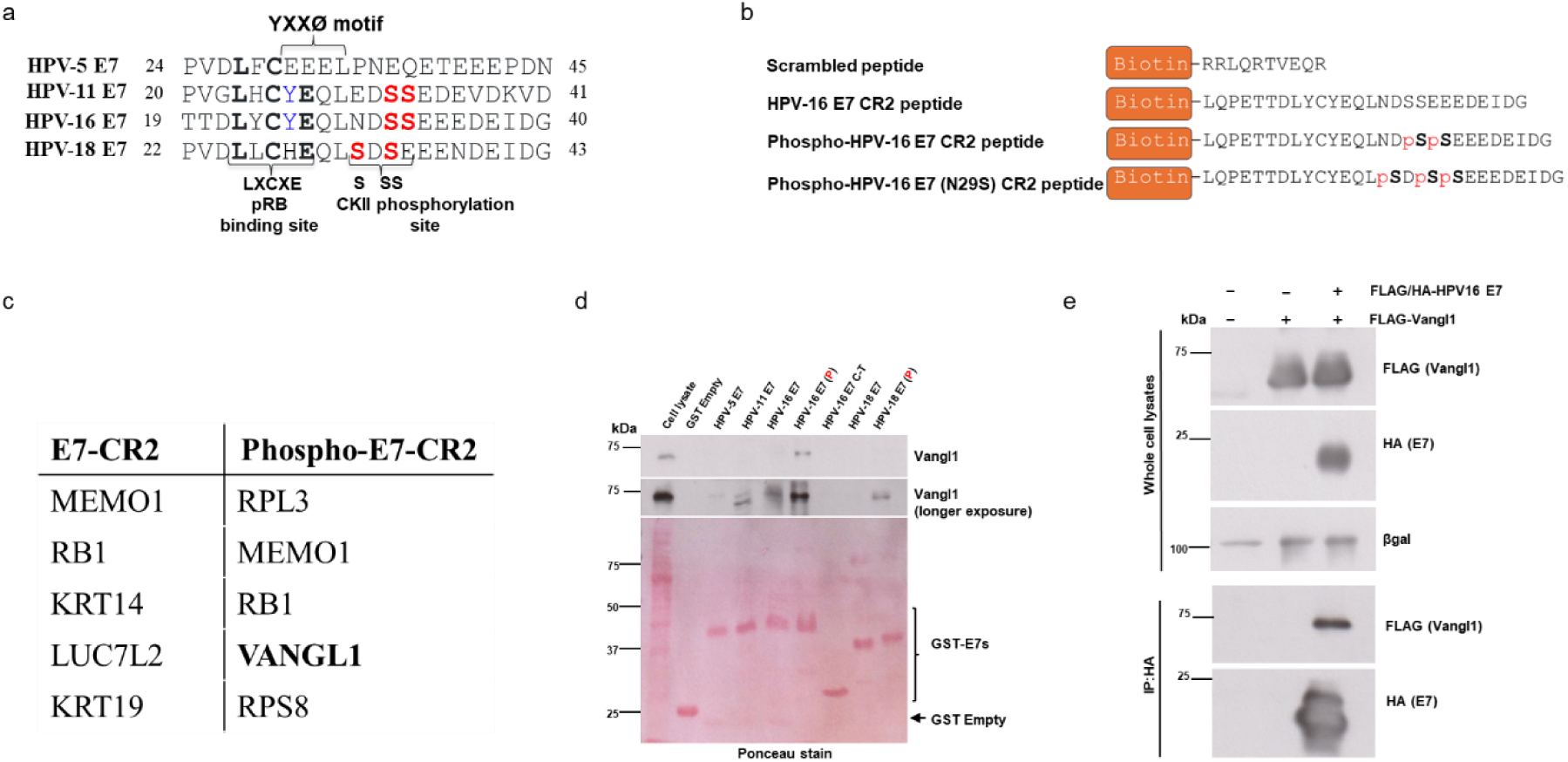
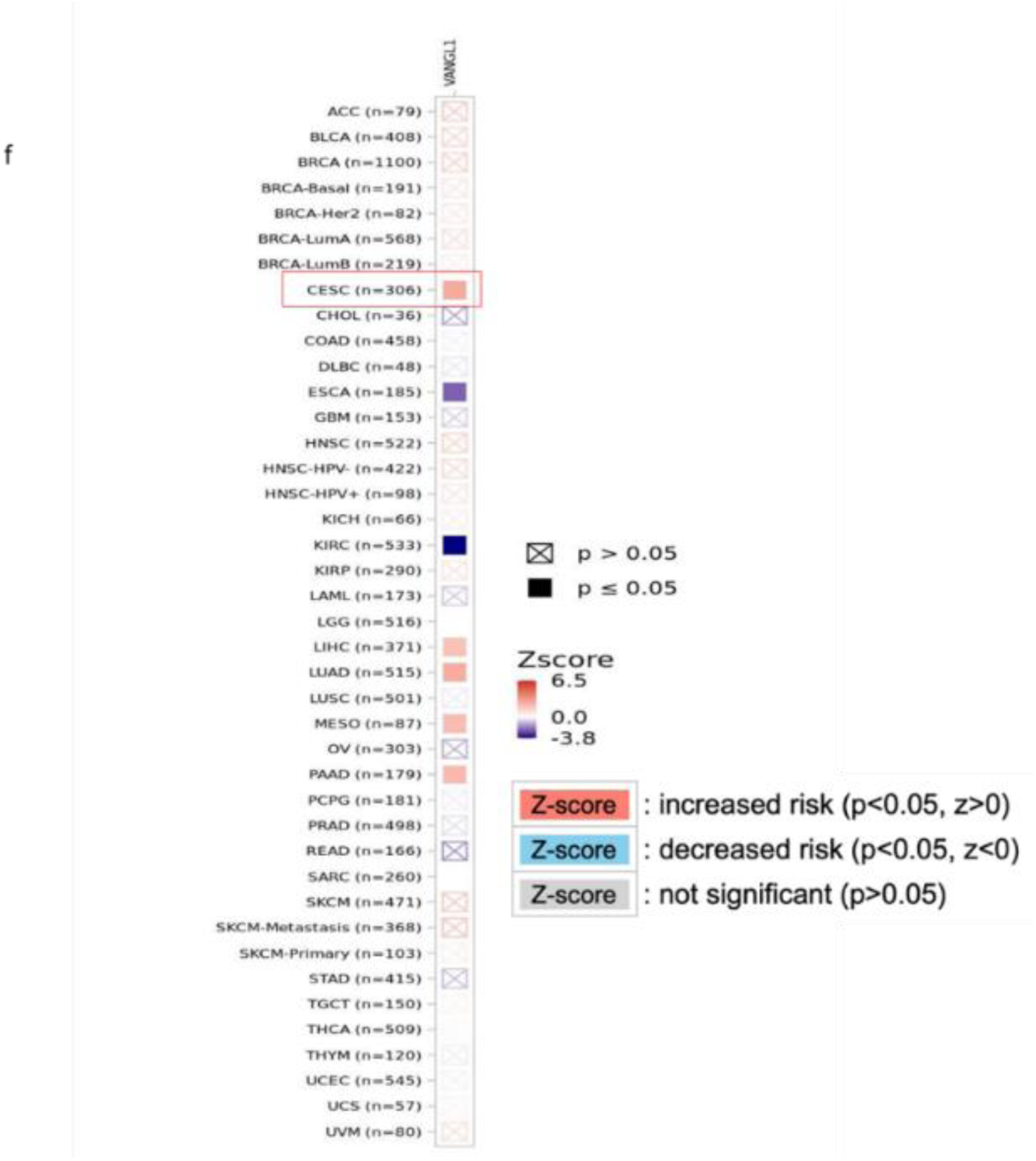
Phosphorylated HPV-16 E7 selectively recruits Vangl1. (a) Schematic of the HPV-16 E7 CR2 region showing the Casein Kinase II (CKII) phosphorylation site at serines 31 and 32/33. (b) Sequences of biotin-tagged peptides corresponding to the HPV-16 E7 CR2 region, synthesized in phosphorylated and non-phosphorylated forms and used as bait in peptide pulldown assays. (c) Mass spectrometry analysis of HaCaT lysates pulled down with phosphorylated or non-phosphorylated E7 peptides. The phosphorylated peptide enriched known E7 interactor pRB and selectively precipitated Vangl1. (d) GST pulldown assays showing strong binding of endogenous Vangl1 to phosphorylated HPV-16 E7, with weak or undetectable binding to HPV-11 E7, non-phosphorylated HPV-16 or HPV-18 E7, phosphorylated HPV-18 E7, or HPV-5 E7. **‘P’** indicates phosphorylated forms of the respective E7s. (e) Co-immunoprecipitation of FLAG-Vangl1 with FLAG/HA-tagged HPV-16 E7 expressed in HEK293 cells confirms the *in vivo* interaction. Beta galactosidase (β-gal) protein expression is used as a loading control in the western blotting analyses. (f) TCGA analysis shows that Vangl1 expression is associated with increased clinical risk in multiple tumour types, including cervical and endocervical cancer (CESC), lower-grade glioma (LGG), lung adenocarcinoma (LUAD), and pancreatic adenocarcinoma (PAAD). The statistical significance is represented as z-score and *p*-value.

Proteomic analysis showed that the canonical E7 interactor, pRB was one of the most enriched proteins, confirming the relevance of the assay. Consistent with previous reports (*52*), the phosphorylated E7 peptide recovered more pRB derived peptides than the non-phosphorylated peptide (Figure 1c), supporting the established role of negative charge at the CKII site in enhancing E7–pRB binding. Of the other proteins precipitated, Vangl1 was notable, for being selectively enriched by the phosphorylated HPV-16 E7 peptide (Figure 1c).

Vangl1 is a core planar cell polarity (PCP) protein that functions as a scaffold in the Wnt/PCP pathway and is essential for cell orientation and migration (*53*). To explore the potential clinical relevance of this interaction, we examined Vangl1 expression in different cancers as published in The Cancer Genome Atlas (TCGA). Elevated Vangl1 gene expression was significantly associated with increased risk in six tumour types, including cervical and endocervical cancer (CESC), lower grade glioma (LGG), lung adenocarcinoma (LUAD), and pancreatic adenocarcinoma (PAAD) (Figure 1f).

We next investigated whether Vangl1 physically interacts with E7 proteins from high-risk (HPV-16, HPV-18), low risk (HPV-11) and cutaneous (HPV-5) HPV types. GST pulldown assays using HaCaT lysates revealed strong binding of endogenous Vangl1 to phosphorylated HPV-16 E7, whereas only weak or undetectable binding was observed with HPV-11 E7, non-phosphorylated HPV-16 or HPV-18 E7, phosphorylated HPV-18 E7, or HPV-5 E7 (Figure 1d). Vangl1 did not bind the HPV-16 E7 C-terminus (residues 41–98), indicating that the N-terminal half mediates this interaction.

To validate the interaction *in vivo*, FLAG-tagged Vangl1 and FLAG/HA-tagged HPV-16 E7 were co-expressed in HEK293 cells. Immunoprecipitation with anti-HA beads confirmed that Vangl1 associates with HPV-16 E7 in cells (Figure 1e). Together, these findings identify Vangl1 as a novel, phosphorylation-dependent interactor of HPV-16 E7.

### CKII phosphorylation allows selective E7–Vangl1 binding

Because Vangl1 and its paralog Vangl2 share 72% identity and overlapping roles in Wnt/PCP signalling (*53*), we examined whether HPV-16 E7 might also interact with Vangl2. Co-immunoprecipitation assays from HEK293 cells expressing FLAG-tagged Vangl1 or Vangl2 with FLAG/HA-tagged HPV-16 E7 showed that E7 binds robustly to Vangl1, but only minimally to Vangl2 (Figure 2a), demonstrating paralog-specific selectivity.

**Figure 2:**
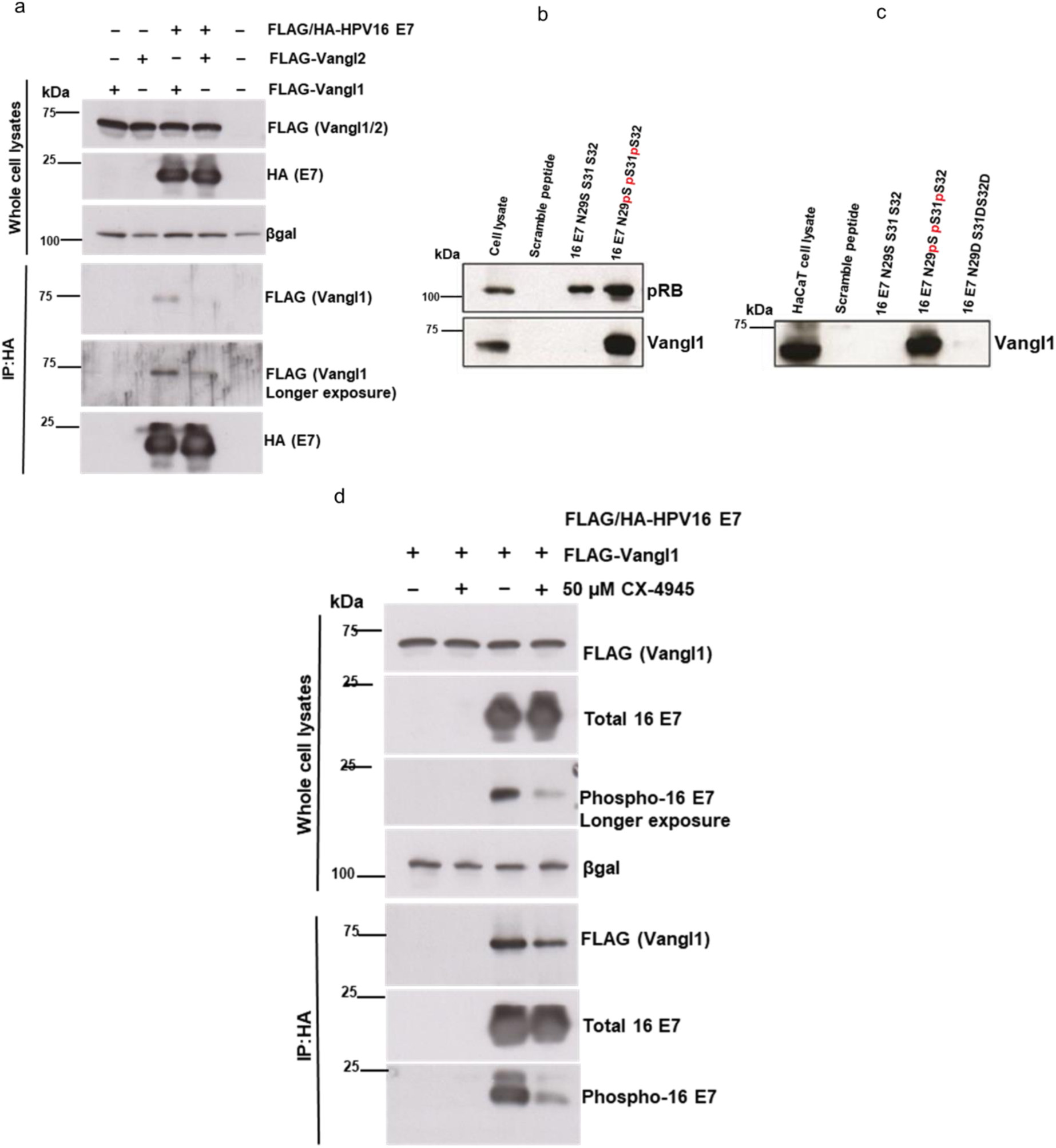
CKII phosphorylation enables HPV-16 E7 to bind specifically to Vangl1. (a) Co-immunoprecipitation of FLAG-tagged Vangl1 or Vangl2 with FLAG/HA-tagged HPV-16 E7 expressed in HEK293 cells. HPV-16 E7 binds strongly to Vangl1 but only minimally to Vangl2. (b) Peptide pulldown assays using phosphorylated and non-phosphorylated HPV-16 E7 N29S CR2 peptides. Vangl1 binds strongly to the phosphorylated N29S peptide but not to its non-phosphorylated form. (c) Pulldown assays using phosphomimic HPV-16 E7 CR2 peptides in which CKII-targeted serines were substituted with aspartic acid. The phosphomimic fails to bind Vangl1. ‘**p’** indicates phosphorylation. (d) Co-immunoprecipitation of FLAG-Vangl1 with FLAG/HA-tagged HPV-16 E7 in HEK293 cells treated with the CKII inhibitor CX-4945. CKII inhibition markedly reduces Vangl1–E7 association. Beta galactosidase (β-gal) protein expression is used as a loading control in the western blotting analyses.

To further assess the role of CKII phosphorylation, we performed peptide pulldown assays using the HPV-16 E7 N29S variant, which undergoes enhanced CKII phosphorylation (*51*). Vangl1 bound strongly to the phosphorylated N29S peptide but not to its non-phosphorylated counterpart (Figure 2b). Notably, the substitution of the CKII-targeted serines with aspartic acid to generate phospho-mimic peptides failed to recapitulate binding (Figure 2c), indicating that negative charge alone is insufficient and that true phosphorylation is required.

We next tested whether CKII activity is required for the interaction *in vivo*. To do this, HEK293 cells co-expressing Vangl1 and HPV-16 E7 were treated with the pharmacological CKII inhibitor CX-4945. CKII inhibition markedly reduced Vangl1 co-immunoprecipitation with E7 (Figure 2d), confirming that CKII-mediated phosphorylation of E7 is essential for Vangl1 binding. Taken together, these results demonstrate that HPV-16 E7 selectively binds Vangl1 in a CK-II phosphorylation-dependent manner.

### HPV E7 expression modulates Vangl1 protein abundance in cervical cancer cells

Having established that Vangl1 is a phosphorylation-specific substrate of E7, we wanted to determine how Vangl1 behaves in HPV-positive cervical cancer cells. For this, we examined Vangl1 protein levels following siRNA-mediated knockdown of E6/E7 or Vangl1 in HPV-positive CaSki and HeLa cells, HPV-negative C-33A cells, and non-transformed HaCaT keratinocytes. It is important to mention that Vangl1 appears as two bands on the blots in most cases – the lower band is unphosphorylated Vangl1 and the upper band is the phosphorylated form.

It can be seen in Figures 3a and b, that the levels of unphosphorylated Vanngl1 (lower band) in siScramble-treated CaSki and HeLa cells are very low but increase upon E6/E7 knockdown in both cell lines increasing total Vangl1 levels relative to the control. It is also notable that, Vangl1 depletion modestly reduced HPV-16 E7 abundance in CaSki cells but had no effect on HPV-18 E7 in HeLa cells, which might reflect the weaker physical interaction between HPV-18 E7 and Vangl1. In contrast, the majority of Vangl1 was phosphorylated in HPV-negative C-33A and HaCaT cells, and its level remained unchanged upon siE6/E7 treatment (Figures 3c, d), confirming that the increase in Vangl1 levels upon E6/E7 knockdown is specific to HPV-positive cervical cancer cells and not an off-target effect of siRNA treatment.

**Figure 3:**
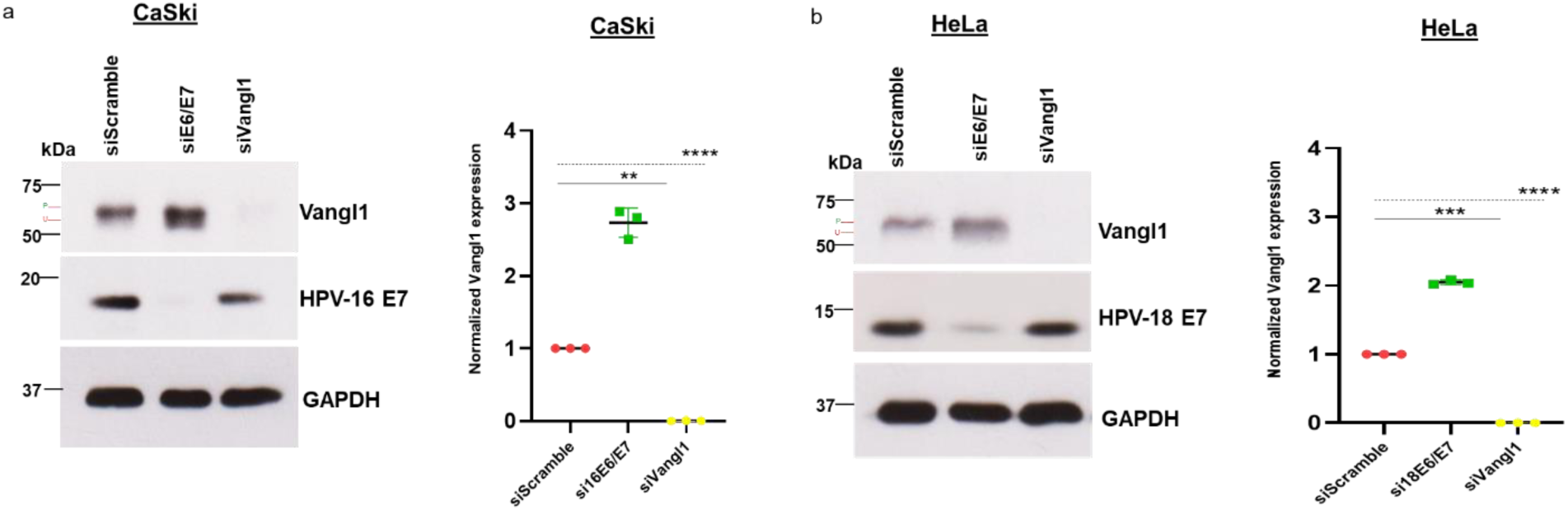
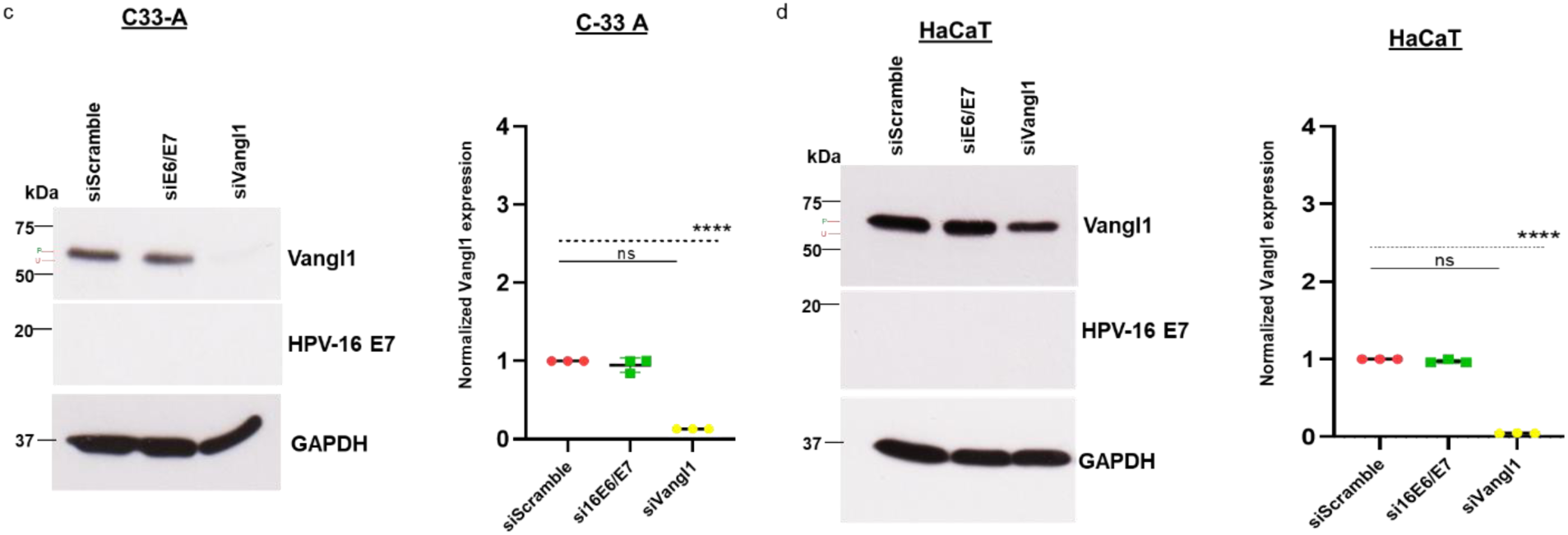
HPV E7 expression modulates Vangl1 protein abundance. Immunoblot analysis of Vangl1 and E7 protein levels following siRNA-mediated knockdown of E6/E7 or Vangl1. (a, b) In HPV-16 E7-positive CaSki cells and HPV-18 E7-positive HeLa cells, E6/E7 depletion increases total Vangl1 abundance. (c, d) In HPV-negative C33A cells and non-transformed HaCaT keratinocytes, Vangl1 remains unchanged following E6/E7 knockdown. GAPDH expression is shown as loading control. Vangl1 band density was normalized to GAPDH band density, and the data was used to plot the densitometry graphs for all cell lines. Data is shown as the fold-changes of normalized Vangl1 levels relative to the control (siScramble) ± standard deviation. *p* values of 3 independent experiments were calculated by Student’s t test. (*p* values of 3 independent experiments were calculated by Student’s t test. (*: *p*-value < 0.05; **: *p*-value <0.01; ***: *p*-value <0.001).

These findings suggest that E7 expression perturbs Vangl1 protein homeostasis in HPV-transformed cervical cancer cells and that Vangl1 may contribute to the stability of HPV-16 E7.

### E7 selectively prolongs half-life of phosphorylated Vangl1

The accumulation of Vangl1 seen in the HPV-positive cervical cancer cell lines prompted us to ask whether E7 might enhance Vangl1’s proteasomal or lysosomal degradation, pathways previously described for its natural turnover (*54*). CaSki cells transfected with either siScramble or siE6/E7 were treated with the proteasome inhibitor CBZ, the lysosome inhibitor chloroquine (CQ), or DMSO. We found that, in control-treated cells, both inhibitors modestly increased Vangl1 levels, with CQ having a stronger effect (Figure 4a). Notably, there was a selective increase in the lower (unphosphorylated) Vangl1 band upon CQ treatment, suggesting that unphosphorylated Vangl1 may be more susceptible to lysosomal degradation, and this is consistent with previous reports (*55*). However, E6/E7 knockdown in CaSki cells increased Vangl1 levels more strongly than inhibitor treatment in siScramble-treated cells. Furthermore, combining E6/E7 knockdown with CQ treatment did not further increase Vangl1 levels, which argues against a model tightly controlled by E7-enhanced lysosomal degradation. These findings suggest that E7 may contribute to the lysosomal degradation of Vangl1, but do not solely explain the accumulation of Vangl1 upon E7 knockdown, suggesting that other mechanisms may also be in play.

**Figure 4:**
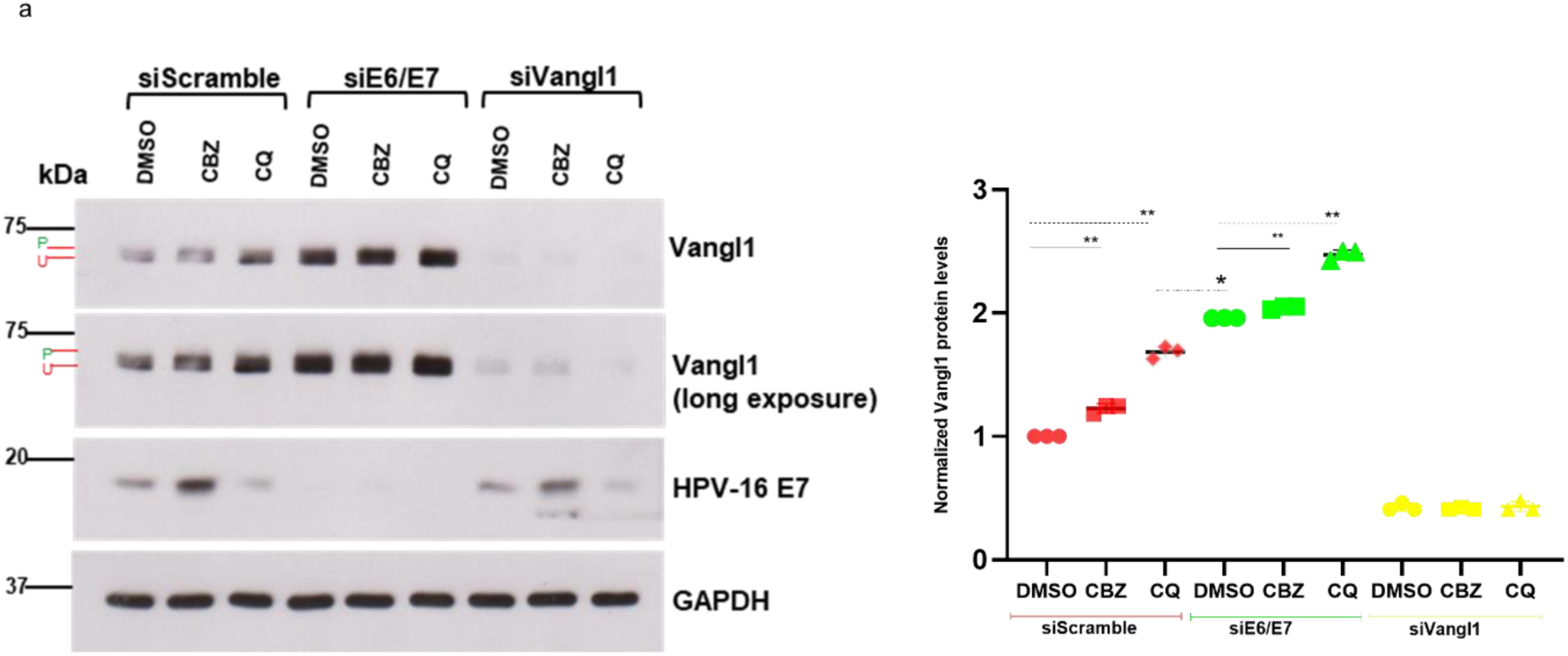
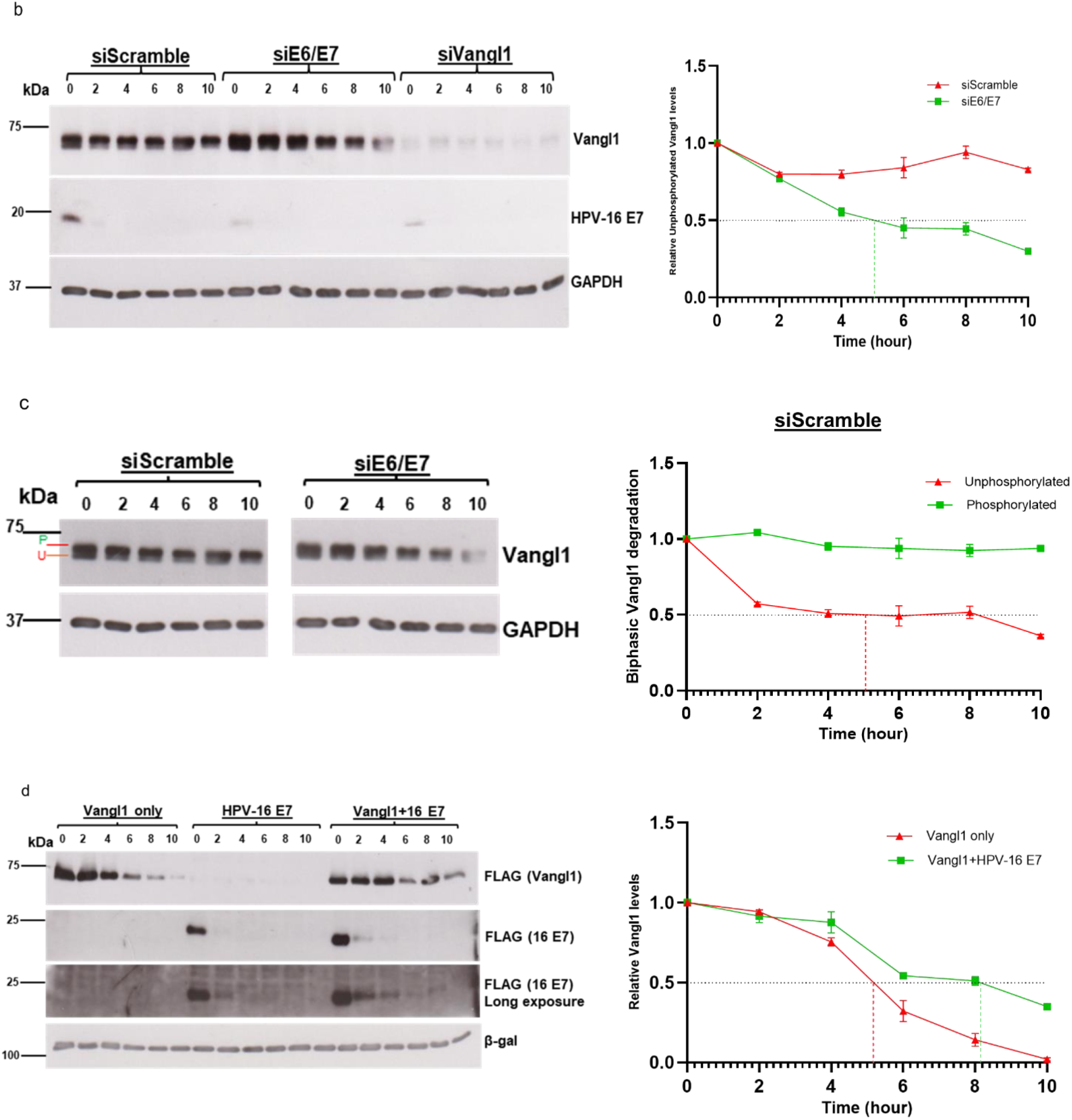
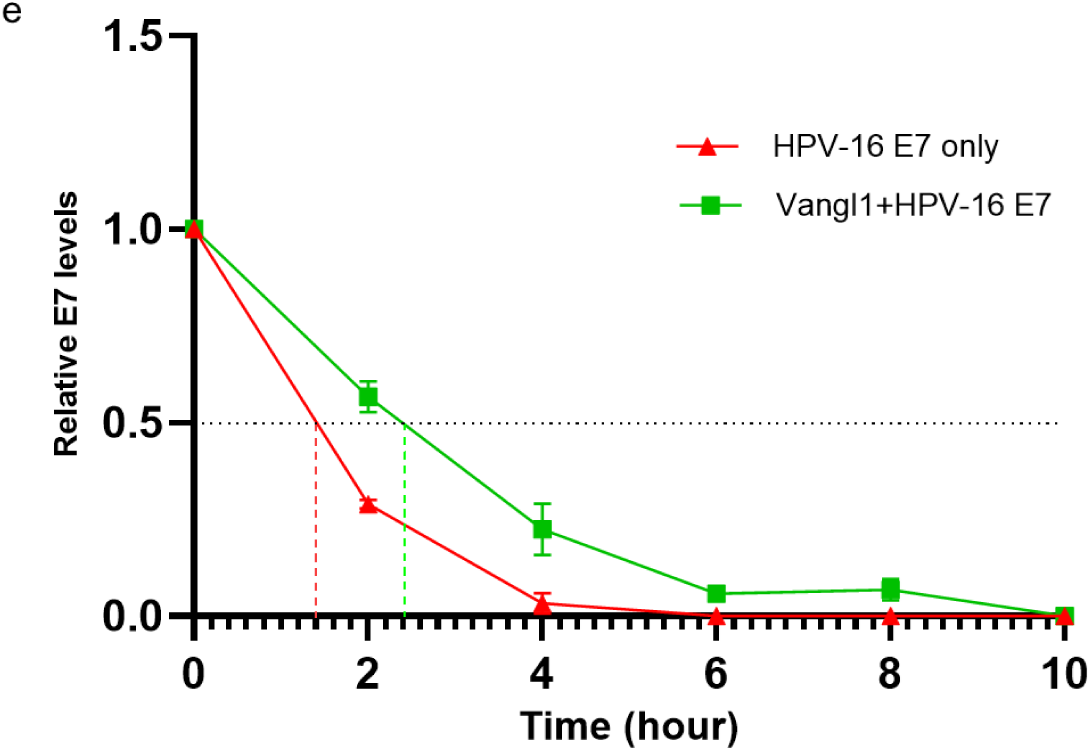
E7 prolongs Vangl1 half-life through a non-canonical stabilization mechanism. (a) Immunoblot analysis of Vangl1 in CaSki cells transfected with control, Vangl1 or E6/E7 siRNAs and treated with the proteasome inhibitor CBZ, the lysosomal inhibitor chloroquine (CQ), or DMSO. E6/E7 knockdown alone increases Vangl1 levels more strongly than either inhibitor. (b) Cycloheximide chase assays in CaSki cells shows that Vangl1 half-life in control cells is longer than in E7-depleted cells. GAPDH expression is shown as loading control. Vangl1 band density was normalized to GAPDH band density, and the data was used to plot densitometry graphs for the cell lines. Data is shown as the fold changes of normalized Vangl1 levels relative to the control (siScramble) ± standard deviation. (c) Long exposure inset of the blot in (b) shows siScramble and siE6/E7 at similar start-point intensities, highlighting biphasic degradation of unphosphorylated and phosphorylated Vangl1 forms in siScramble cells. (d) Cycloheximide chase assays in HEK293 cells overexpressing Vangl1 with or without HPV-16 E7. Vangl1 half-life increases when co-expressed with E7. (e) Densitometry showing reciprocal stabilization of E7 by Vangl1. E7 half-life increases when co-expressed with Vangl1. Dotted lines on time-kinetic graphs indicate specific half-life. β-gal expression is shown as loading control. Vangl1 band density was normalized to the β-gal band density, and the data was used to plot a densitometry graph. Data is shown as the fold changes of normalized Vangl1 levels relative to the control (Vangl1 only) ± standard deviation. *p* values of 3 independent experiments were calculated by Student’s t test. (*p* values of 3 independent experiments were calculated by Student’s t test. (*: *p*-value < 0.05; **: *p*-value <0.01; ***: *p*-value <0.001).

To investigate this, we performed Cycloheximide chase assays to monitor the posttranslational stability of Vangl1 in the presence of E7. In control-treated CaSki cells, the half-life of total Vangl1 exceeded 10-h but decreased to 8.4-h upon E6/E7 knockdown (Figure 4b). This was intriguing because the pool of total Vangl1 in E6/E7-depleted CaSki cells is at least twice that of control-treated cells across different experiments. Strikingly, the control-treated cells exhibited a biphasic degradation pattern, unlike the siE6/E7-treated cells where steady concurrent degradation was seen. In Figure 4c, we show that unphosphorylated Vangl1 (lower band labelled ‘U’) was initially degraded rapidly, reaching its half-life in the first 5-h and then remained stable until the 8^th^ hour followed by a decline. This aligns with our earlier observation that CQ treatment selectively rescued unphosphorylated Vangl1 in siScramble-treated cells. In contrast, the half-life of the phosphorylated form (upper band labelled ‘P’) exceeded 10-h, much longer than the reported 3-6-h in normal cells (*54*), indicating that phosphorylated Vangl1 undergoes aberrant proteostasis in HPV-transformed cervical cancer cells. Together, these observations indicate that the increase in total Vangl1 pool following E7 loss is not the result of enhanced stability *per se*, but instead reflects a non-canonical regulatory mechanism.

To validate this, we overexpressed Vangl1 with or without HPV-16 E7 in HEK293 cells. Vangl1 half-life increased from 5.2 h to 8.2 h when co-expressed with E7 (Figure 4d). Conversely, E7 half-life increased from 1.2 h to 2.2 h when co-expressed with Vangl1 (Figure 4d, e). In addition to the difference in half-life, E7 remained detectable 8 h post cycloheximide treatment when co-expressed with Vangl1 while it was no longer visible after the 2^nd^ hour of treatment when expressed alone (Figures 4c, d). Taken together, these results reveal a reciprocal stabilization mechanism in which E7 aberrantly prolongs the half-life of phosphorylated Vangl1 for its stabilization.

### E7 alters Vangl1 localization in cervical cancer cells

Because protein sequestration, aggregation and mislocalization are established mechanisms by which protein turnover and overall protein abundance are altered (*56, 57*), we examined whether E7 might affect Vangl1 subcellular distribution within cervical cancer cells. Subcellular fractionation of CaSki cells showed that no Vangl1 was present in the cytoskeletal fraction in control cells, but that phosphorylated Vangl1 relocates there upon E6/E7 knockdown (Figure 5a). In addition, there is an increase in the levels of both unphosphorylated Vangl1 and phosphorylated Vangl1 pools in the cytoplasmic fraction upon E6/E7 knockdown, whereas membrane-associated Vangl1 remained largely unchanged. These findings suggest that Vangl1 trafficking and metabolism is impaired in the presence of E6/E7, reflected by limited export or enhanced cytoplasmic instability, prolonged retention at the membrane, and no localization to the cytoskeletal regions.

**Figure 5:**
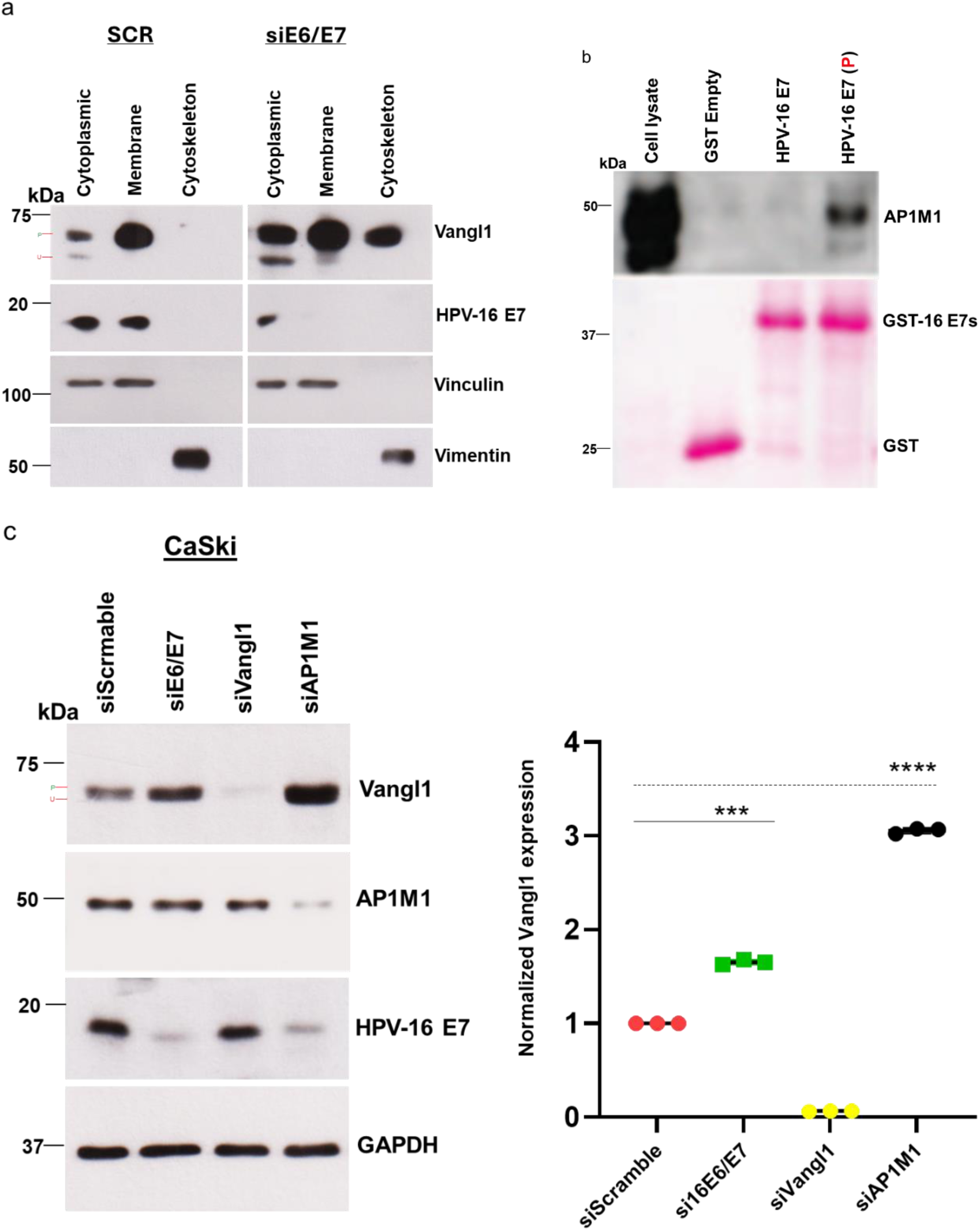
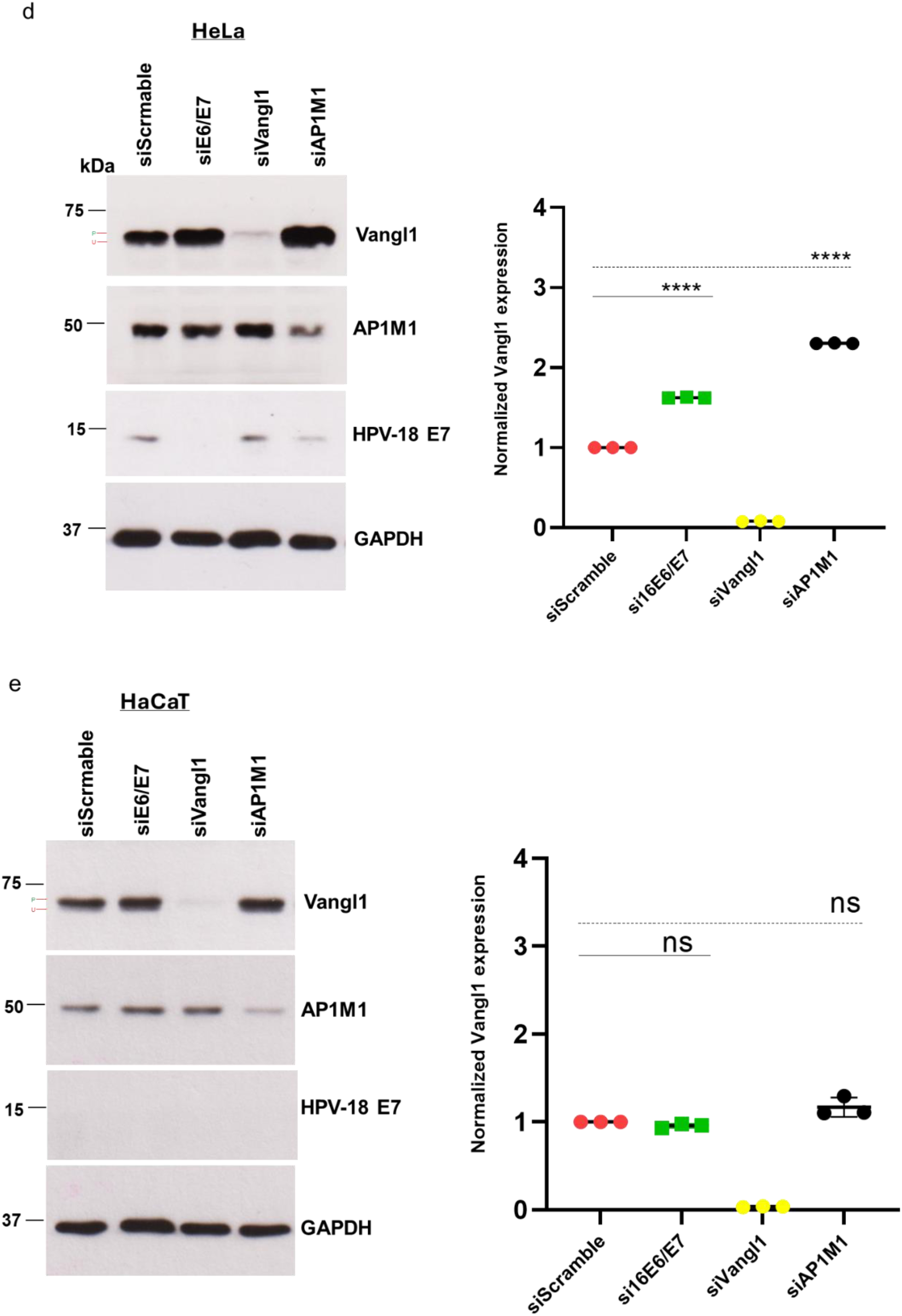
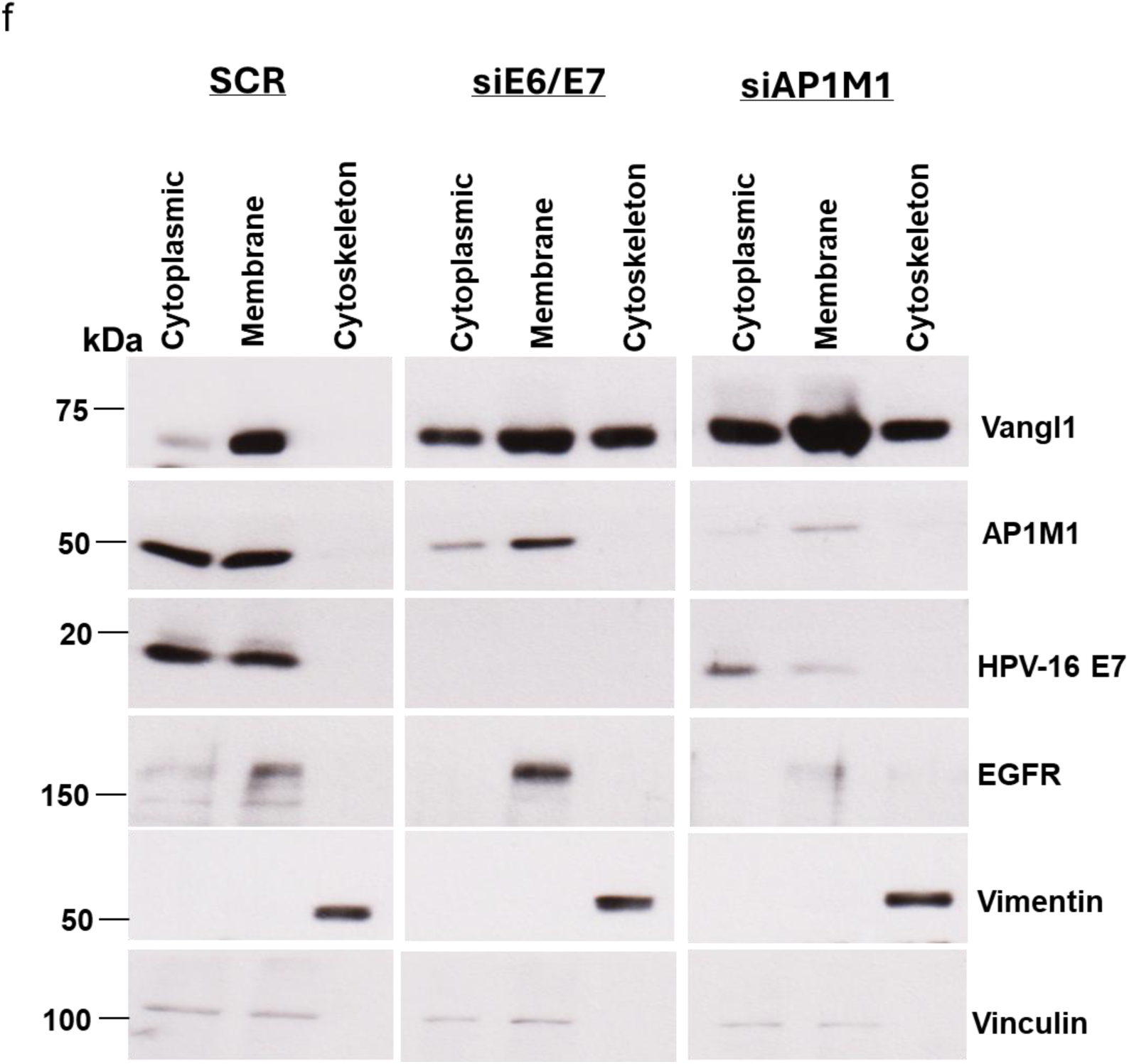
E7 remodels AP1M1-dependent trafficking to alter Vangl1 localization. (a) Subcellular fractionation of CaSki cells transfected with control or E6/E7 siRNAs. Vangl1 is absent from the cytoskeletal fraction in control cells, but becomes detectable upon E6/E7 knockdown. (b) GST pulldown assays showing that AP1M1 selectively binds phosphorylated HPV-16 E7. (c–d) Immunoblot analysis of CaSki and HeLa cells shows that Vangl1 abundance increases upon E6/E7 depletion and is further increased by AP1M1 knockdown. (e) Immunoblot analysis of HaCaT keratinocytes shows that neither E6/E7 knockdown nor AP1M1 knockdown results in increased Vangl1 levels. (f) Subcellular fractionation of CaSki cells after AP1M1 knockdown shows Vangl1 accumulation across cytoplasmic, membrane, and cytoskeletal compartments mirroring E6/E7 knockdown. GAPDH expression is shown as loading control. Vangl1 band density was normalized to GAPDH band density, and the data was used to plot densitometry graphs for the cell lines. Data is shown as the fold changes of normalized Vangl1 levels relative to the control (siScramble) ± standard deviation. *p* values of 3 independent experiments were calculated by Student’s t test. (*p* values of 3 independent experiments were calculated by Student’s t test. (*: *p*-value < 0.05; **: *p*-value <0.01; ***: *p*-value <0.001).

Vangl1 trafficking is regulated by the μ1 subunit of the AP-1 clathrin adaptor complex (AP1M1), which mediates sorting between the trans-Golgi network, plasma membrane, cytoskeleton and lysosomes (*58–60*). Given AP1M1’s central role in Vangl1 trafficking, we asked whether AP1M1 might participate in the E7-driven regulation of Vangl1. First, we used GST pulldown assays to determine whether Vangl1 interacts physically with AP1M1. These showed that phosphorylated, but not unphosphorylated HPV-16 E7 binds to AP1M1 (Figure 5b), mirroring the Vangl1 interaction pattern.

We then wanted to assess the dynamics of Vangl1, AP1M1, and E7 interactions in cervical cancer cell lines and in normal keratinocytes. We had found earlier (Figure 3), that E6/E7 knockdown increases Vangl1 abundance in whole cell lysates of CaSki and HeLa cells; and it can be seen in Figures 5c and d that, this effect was even more pronounced when AP1M1 is also depleted. Notably, AP1M1 knockdown also led to a substantial reduction in E7 levels in both cell lines, indicating that HPV-18 E7 uses a similar mechanism as HPV-16 E7, to modulate Vangl1, despite its weaker physical interaction. Collectively, these findings suggest; first, that E7 probably sequesters AP1M1, altering the sorting and localization of Vangl1, such that loss of E7 restores proper trafficking; and second, that depletion of AP1M1 destabilizes E7, effectively shutting down the impaired trafficking and its associated cytoplasmic instability, which in turn reduces Vangl1 turnover. Both suggestions point to AP1M1 as a contributing factor, enabling E7 to manipulate Vangl1 homeostasis. In contrast, neither E6/E7 knockdown (as control) nor AP1M1 knockdown led to an increase in Vangl1 in HaCaT keratinocytes (Figure 5e), indicating that this regulatory axis is specific to HPV-transformed cervical cancer cells.

Further delineating this relationship, subcellular fractionation of CaSki cells following AP1M1 knockdown revealed Vangl1 accumulation across cytoplasmic, membrane, and cytoskeletal fractions (Figure 5f), closely resembling the pattern observed with E6/E7 knockdown. Collectively, these findings indicate that E7 expression negatively regulates Vangl1 sorting and trafficking in HPV-transformed cervical cancer cells.

### Vangl1 depletion disrupts 3D spheroid integrity and mirrors E6/E7 loss in cervical cancer cells

To visualize the spatial dynamics of Vangl1 in the presence of HPV-16 E7 in light of our findings in Figure 5, we performed immunofluorescence staining in HEK293 cells expressing Vangl1 alone, HPV-16 E7 alone, or both proteins together. As shown in Figure 6a, cells expressing Vangl1 alone exhibit a structured, peripheral distribution of Vangl1 (red fluorescence of Vangl1) consistent with membrane and cytoskeletal localization. In contrast, co-expression of Vangl1 and HPV-16 E7 (green fluorescence) markedly altered Vangl1 localization. Vangl1 becomes more diffuse and cytoplasmic, with reduced levels in the cortex. E7 expressed alone appears to be predominantly nuclear, but with visible cytoplasmic pools that become more concentrated upon co-expression with Vangl1. Strikingly, some cells (red arrows) show a redistribution of E7 from the nucleus to the peripheral compartments, where it co-localizes (regions of yellow fluorescence) with Vangl1. The merged and zoomed panels highlight the co-localization of Vangl1 and E7 in intracellular compartments. These observations further support the hypothesis that HPV-16 E7 hijacks Vangl1, leading to its mislocalization, away from the cell cortex and into compartments where E7 localizes. These data, together with the biochemical assays, suggest a model in which Vangl1 is sequestered in cervical cancer cells, impairing its scaffold function and contributing to its defective turnover.

**Figure 6:**
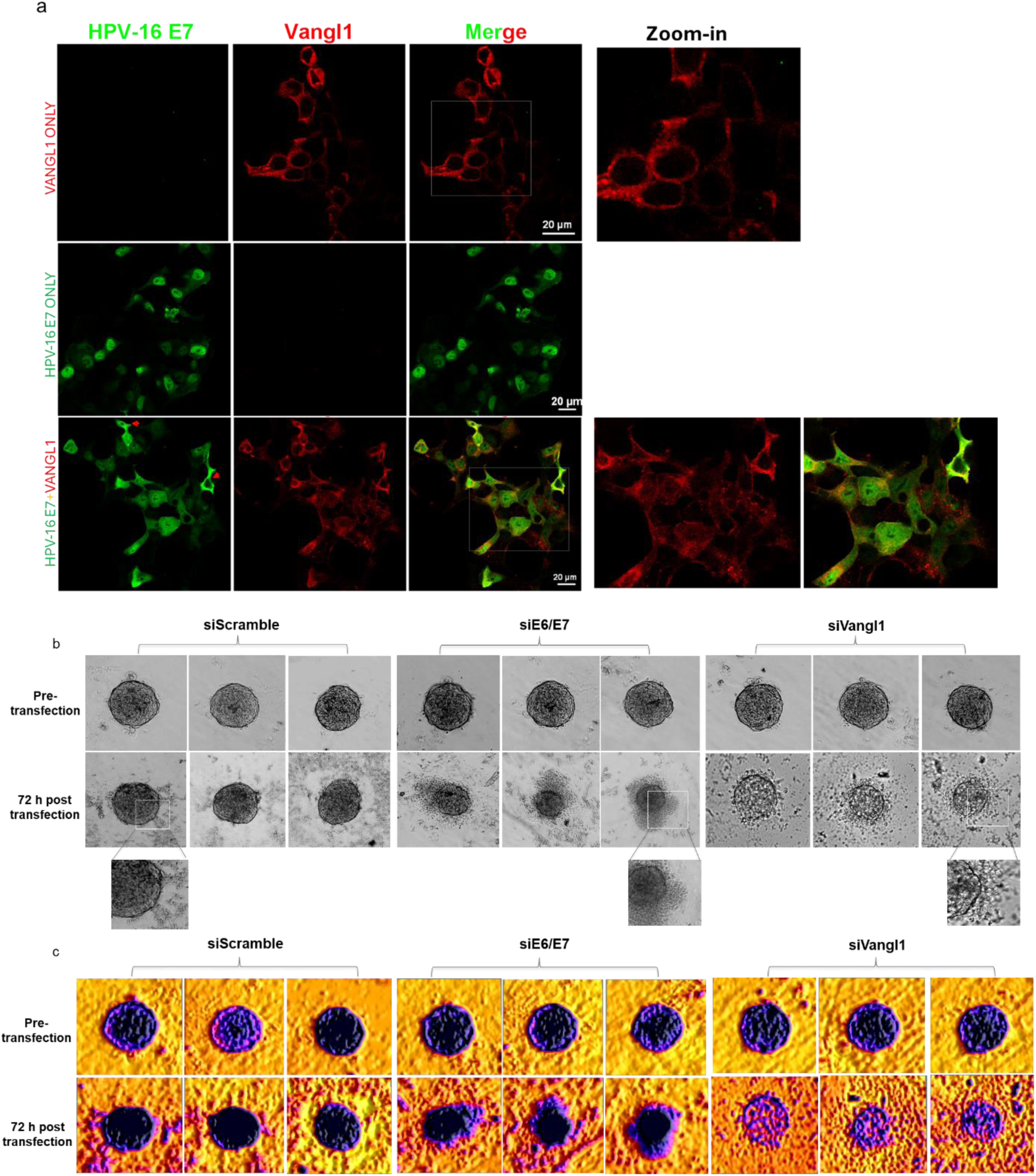
Vangl1 depletion disrupts 3D spheroid integrity. (a) Confocal microscopy of HEK293 cells expressing Vangl1 alone, HPV-16 E7 alone, or both proteins together, show diffuse, cytoplasmic Vangl1 with reduced cortical levels in cells expressing E7. Merged and zoomed images show co-localization of Vangl1 and E7 within intracellular compartments. (b) Brightfield images of CaSki spheroids following siRNA-mediated knockdown of E6/E7 or Vangl1 showing spheroids with pronounced structural disintegration, irregular edges, and reduced cohesion. (c) 3D surface plots of spheroids highlight architectural defects. Control spheroids show uniform signal intensity and well-defined boundaries, while E6/E7 and Vangl1 knockdown spheroids display diffuse signal, fragmented contours, and disrupted core organization. Micrographs were prepared using the Quickfigures plugin of ImageJ. The brightness of the panels was uniformly enhanced post image processing.

We then wanted to know how this mislocalization might impact Vangl1 function in a physiologically relevant context (*61*). To do this, we examined Vangl1’s role in cervical cancer spheroids, where the 3D architecture more faithfully preserves the polarity cues. For polarity proteins, mislocalization can have significant implications, including disruption of normal cell architecture, signalling, and tissue organization, potentially contributing to pathological states (*62–64*). To assess these functional consequences, we examined spheroid morphology in CaSki cells following siRNA-mediated knockdown of E6/E7 or Vangl1. Brightfield imaging revealed that control spheroids (siScramble) maintained their compact, rounded architecture over 72 h, indicative of intact polarity and adhesion signalling (Figure 6b). In contrast, spheroids subjected to E6/E7 or Vangl1 depletion exhibited marked structural disintegration, with irregular edges and loss of cohesion, that are more striking in siE6/E7-depleted spheroids. The 3D surface plots further highlighted the uniform signal intensity and well-defined boundaries of control spheroids, whereas the E6/E7 and Vangl1 knockdown spheroids showed diffuse signals, fragmented contours, and disrupted core organization (Figure 6c). The phenotype of the siE6/E7-treated spheroids indicate that the continuous expression of E6/E7 is crucial for the viability and structural maintenance of the spheroids. Furthermore, the parallel phenotype of siVangl1-treated spheroids suggest that Vangl1 is a critical downstream effector of E7 in maintaining essential epithelial architecture and integrity. Thus, the spheroid models confirm that E7-driven Vangl1 mislocalization is essential for preserving tissue-like organization in transformed cervical cancer cells.

### Vangl1 supports collective invasion and matrix-responsive architecture in transformed cervical cancer cells

Having found that Vangl1 is important for maintaining epithelial architecture in CaSki spheroids, we next investigated the implication of this structural role. The Wnt/PCP pathway is well established as a regulator of cell migration and apical-basal coordination (*65*), and Vangl dysregulation has been linked to metastatic progression (*66–68*). We therefore asked whether Vangl1 could contribute to invasive behaviour in transformed cervical cancer cells. In a 2D Matrigel invasion assay, siRNA-mediated depletion of Vangl1 led to a marked reduction in the number of invading cells, compared with siScramble controls, which displayed dense invasion through the matrix (Figure 7a). Quantification confirmed a statistically significant decrease in invasion upon Vangl1 knockdown, indicating that Vangl1 is required for efficient invasion in this context (Figure 7b).

**Figure 7:**
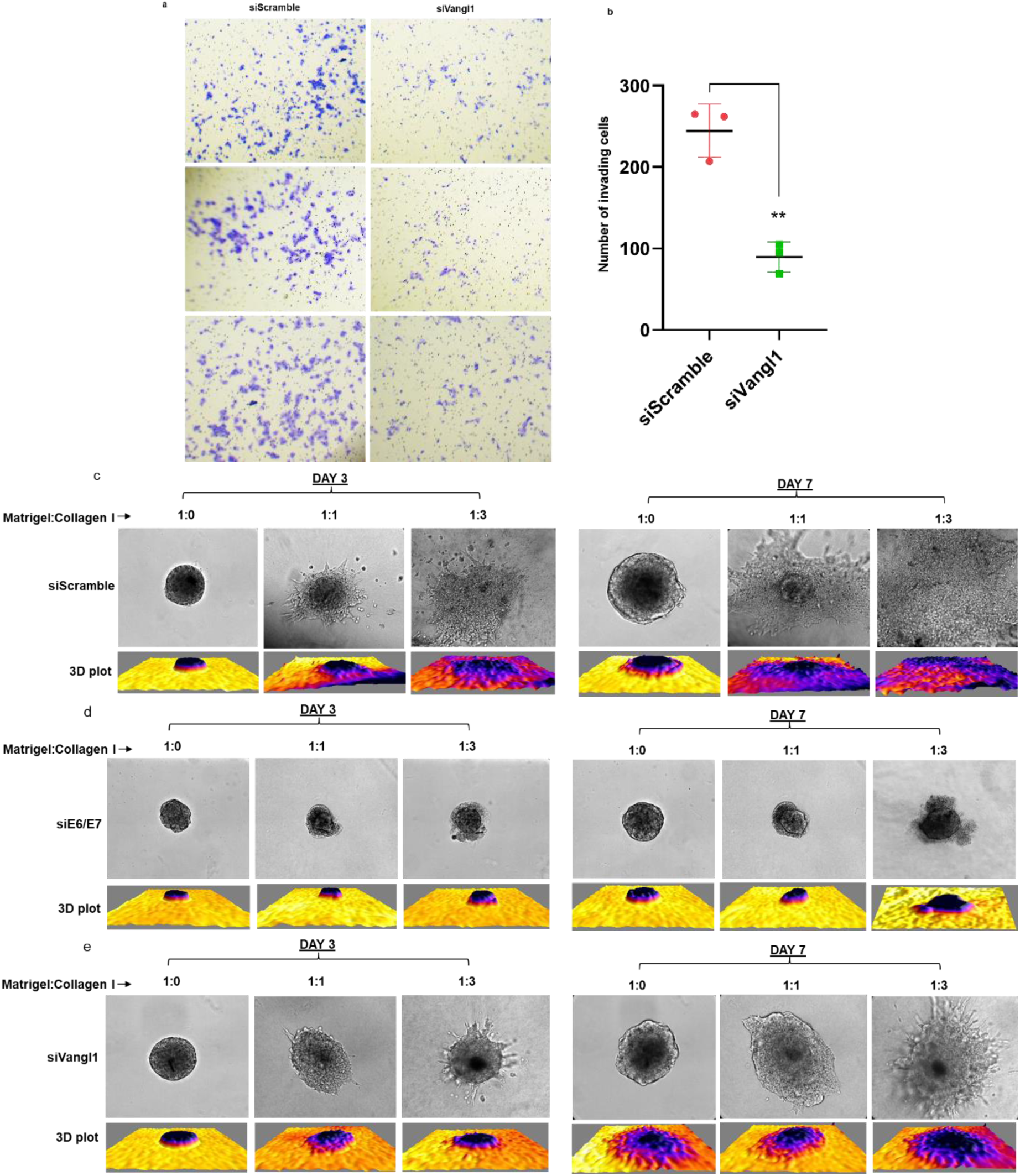
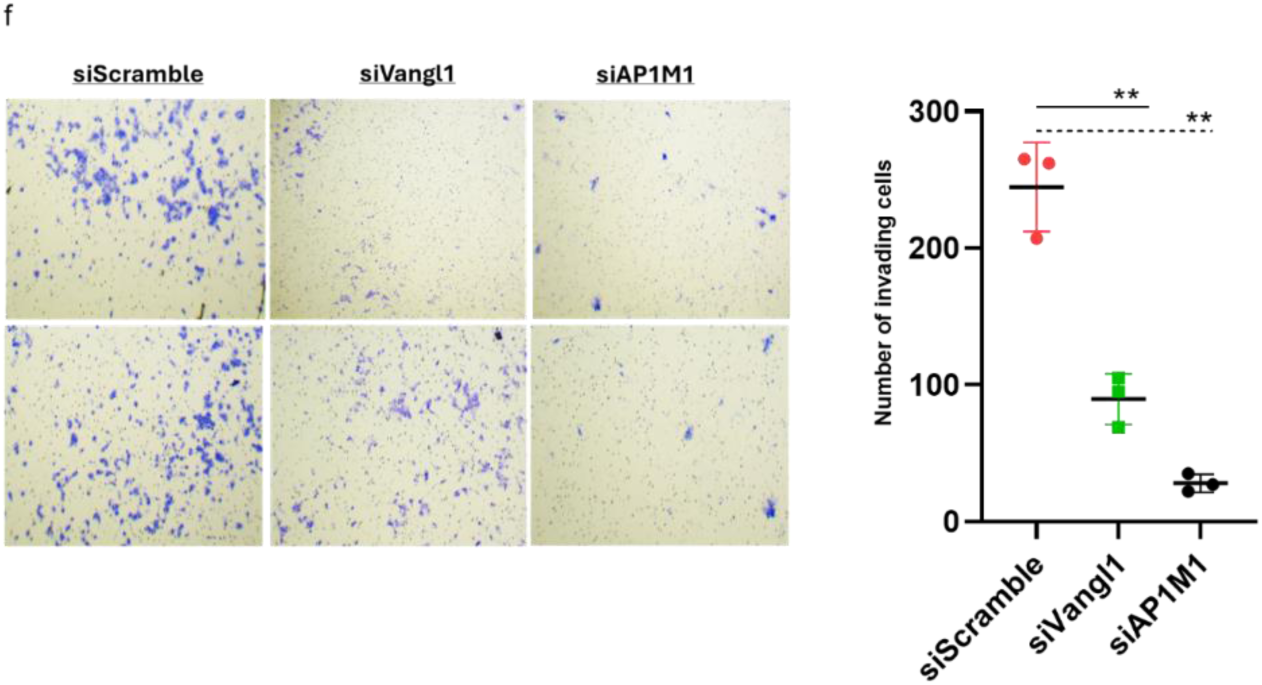
Vangl1 is essential for maintaining invasive phenotype in cervical cancer cells. (a) Brightfield representative images from 2D Matrigel invasion assays in CaSki cells show dense invasion in control cells, whereas Vangl1-depleted cells show markedly reduced invasion. (b) Quantification of invading cells using the ImageJ Cell counter tool confirms a significant decrease in invasion upon Vangl1 knockdown. (c) Brightfield images of spheroid invasion across increasing collagen content (Matrigel:Collagen 1:1 and 1:3) show collective invasion in control spheroids; (d) E6/E7-depleted spheroids fail to generate organized invasive protrusions in either matrix condition; (e) Vangl1-depleted spheroids similarly lack robust invasive structures and display matrix-dependent architectural defects. Corresponding 3D surface plots below the micrographs reflect spheroidal architectural integrity. Invasiveness of the spheroids was monitored for 7 days and imaged at 20x magnification. (f) Brightfield representative images from 2D Matrigel invasion assays following AP1M1 knockdown reveal a pronounced reduction in invasion. Invading cells in the 2D assays were stained with crystal violet and imaged using a brightfield microscope at 10x magnification. *p* values of 3 independent experiments were calculated by Student’s t test. (*: *p*-value < 0.05; **: *p*-value <0.01; ***: *p*-value <0.001; ****: *p*-value <0.0001). Micrographs were prepared using the Quickfigures plugin of ImageJ. The brightness of the panels was uniformly enhanced post-image processing.

To further assess the role of Vangl1 in a physiologically relevant 3D environment, we examined spheroid behaviour across increasing collagen content. Control spheroids (siScramble) exhibited clear invasive protrusions, particularly in the 1:1 Matrigel:Collagen matrix at Day 3, where cohesive, finger-like extensions radiated from an intact spheroid core – hallmarks of organized collective invasion (*69, 70*) (Figure 7c). These protrusions expanded by Day 7, accompanied by sheet-like spreading highlighted by intensely diffused colour change in the 3D surface plots. In the 1:3 matrix, control spheroids underwent complete dissolution of the core and displayed pronounced sheet-like invasion at both timepoints, suggesting that increased matrix stiffness enhances the aggressiveness and collective invasiveness of CaSki cells.

In contrast, neither E6/E7-depleted nor Vangl1-depleted spheroids generated comparable protrusive structures under any matrix condition. E6/E7 knockdown caused early loss of cohesion, edge-fraying, and progressive fragmentation (Figure 7d). Similarly, Vangl1-depleted spheroids displayed significantly fewer invasive protrusions, with the most visible extensions occurring in the 1:3 matrix (Figure 7e). In addition to the loss of invasiveness, Vangl1 depletion produced distinct, matrix-dependent structural defects. In the 1:1 matrix, where control spheroids initiated collective invasion, siVangl1 spheroids instead showed shape distension and short, blunt extensions indicative of compromised epithelial organization rather than directed invasion. In the stiffer 1:3 matrix, the spheroids appeared more compact but exhibited scalloped edges and a striking mono-directional invasion rather than the radial features seen in controls. These suggest that the underlying architectural instability persists but is partially masked by the mechanical constraints of the collagen-rich environment.

Having established that Vangl1 is required for efficient invasion, we next examined whether AP1M1 likewise contributes to the invasive behaviour of transformed cervical cancer cells. Using the 2D Matrigel invasion assay, siScramble cells displayed robust invasion, as previously seen (Figure 7f). Also consistent with our earlier findings is that Vangl1 depletion markedly reduced invasion, resulting in sparse distributions of invading cells. Notably, AP1M1 knockdown produced a similarly pronounced reduction in invasion, with even lower cell numbers than those observed in the siVangl1 condition. This phenotypic overlap indicates that the E7-AP1M1-Vangl1 regulatory relationship is essential for invasive capacity in transformed cervical cancer cells.

Collectively, these observations demonstrate that E7 modulation of Vangl1 is required to maintain the invasive behaviour of transformed cervical cancer cells across diverse extracellular matrix contexts.

### Vangl1 enrichment at the leading edge of migrating cells supports collective invasion

Having established that Vangl1 is essential for collective invasion, we then examined whether it contributes to the polarity cues that support this invasion phenotype. We examined Vangl1 subcellular localization by immunofluorescence in invading spheroids embedded in 1:1 Matrigel:Collagen matrix after 24 h. In siScramble spheroids, Vangl1 displayed strong enrichment with a striking bias toward the outer edge of leading cells (orange arrows), forming continuous arcs at the migration front (Figure 8a). This localization coincided with elongated, directionally oriented cell shapes, indicating a role of Vangl1 in establishing the front–rear cell coordination required for collective movement. In addition, single disseminated cells (white arrow) were also seen, suggesting a multimodal invasive capacity of the spheroids. In contrast, siVangl1 spheroids showed markedly reduced Vangl1 signal, with no detectable enrichment at leading edges, accompanied by irregular spheroidal morphology (red arrows), and loss of clear directional orientation (Figure 8b). Previous reports have shown that the localization of polarity proteins to the edge of invading cells is highly significant (*68, 71*); it marks the conversion of a static, apico-basal polarized epithelial cell into a motile, front-rear polarized cell capable of movement and invasion. Therefore, these findings indicate that Vangl1 is required not only for maintaining epithelial architecture, but also for positioning polarity machinery at the migration front to allow cohesive, collective invasion of transformed cervical cancer cells.

**Figure 8:**
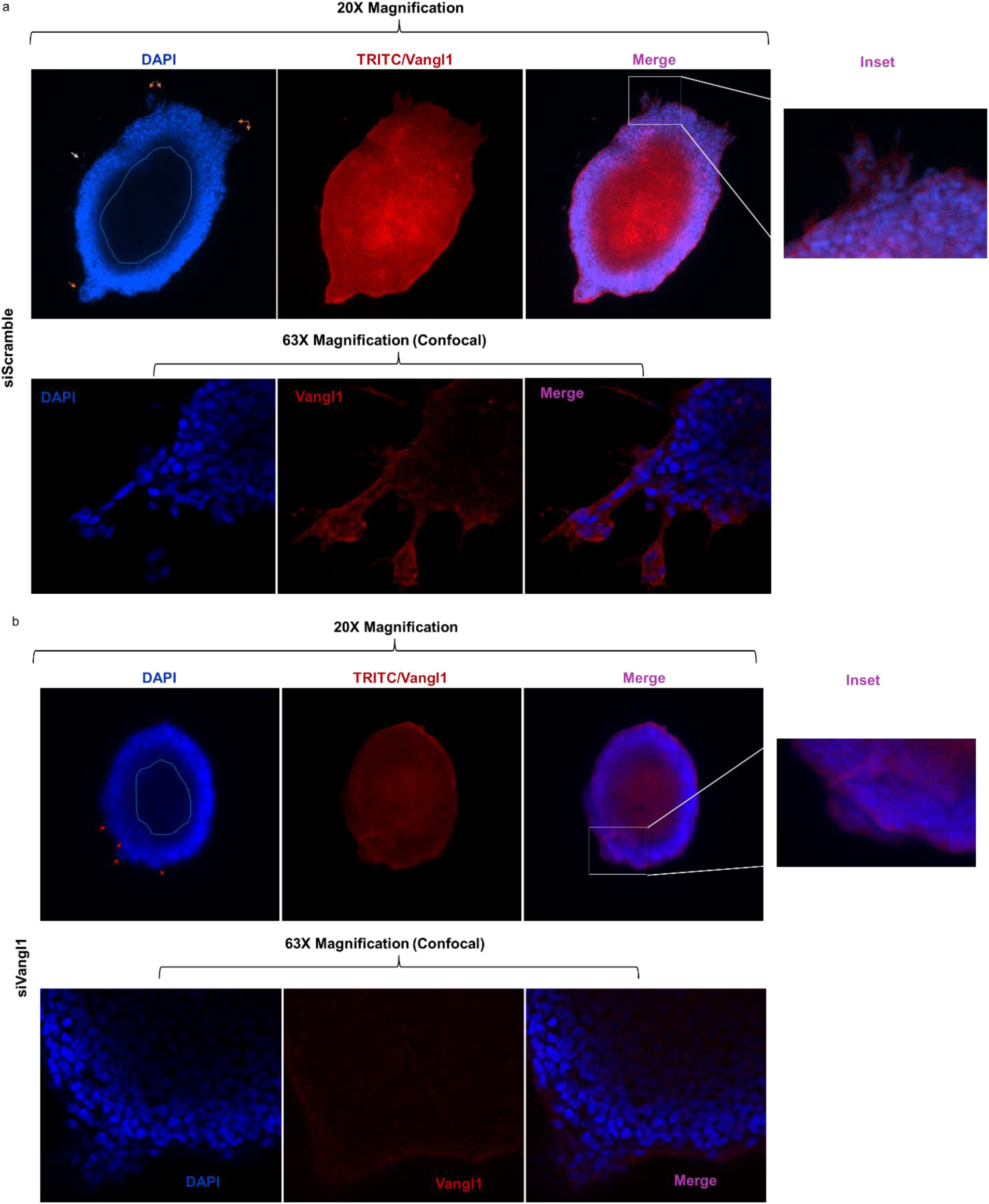
Vangl1 is enriched at the leading-edge of invading protrusions of CaSki spheroids. (a) Top panel shows fluorescent images of invading control siScramble CaSki spheroids in 1:1 Matrigel:Collagen matrix at 20x magnification. The bottom panel shows immunofluorescence images showing a section of the invading protrusions displaying strong Vangl1 enrichment at the outer edge of leading cells, visualized by confocal microscopy at 63x magnification. (b) siVangl1 spheroids show markedly reduced Vangl1 signal with no detectable leading-edge enrichment. Loss of Vangl1 results in irregular spheroid morphology and scalloped edges (red arrows). Cell nuclei were stained with Hoeschst 33342, and endogenous anti-Vangl1 antibody was counterstained with anti-mouse Alexa Fluor 594. Micrographs were prepared using the Quickfigures plugin of ImageJ. The brightness of the panels was uniformly enhanced post image processing.

### Vangl1 knockdown enhances chemosensitivity of cervical cancer spheroids

Based on the established oncogenic contributions of Vangl1, we wanted to evaluate how Vangl1 influences chemosensitivity in CaSki spheroids. Forty-eight hours (48-h) after siRNA transfection (recorded as 0-h of treatment), we monitored spheroid morphology over the subsequent 48-h, following treatment with DMSO, Palbociclib, Cisplatin, or Etoposide. Under DMSO, siScramble spheroids remained compact and cohesive, while siE6/E7 and siVangl1 spheroids exhibited progressively compromised architecture, including edge-fraying and reduced height, indicating a baseline structural instability (Figure 9a). Palbociclib treatment induced mild disintegration in siScramble spheroids, but triggered pronounced dissociation in E6/E7- and Vangl1-depleted spheroids (Figure 9b). Cisplatin (Figure 9c) and Etoposide (Figure 9d) treatments also replicated these effects: siScramble spheroids showed moderate architectural disruption, whereas siE6/E7 and siVangl1 spheroids exhibited fragmented contours, loss of central density, significantly dissociated cells, reduced size and a halo of cell debris. While Palbociclib largely disrupted the compactness of the spheroids, Cisplatin and Etoposide caused peripheral disintegration and reduction in spheroid sizes.

**Figure 9:**
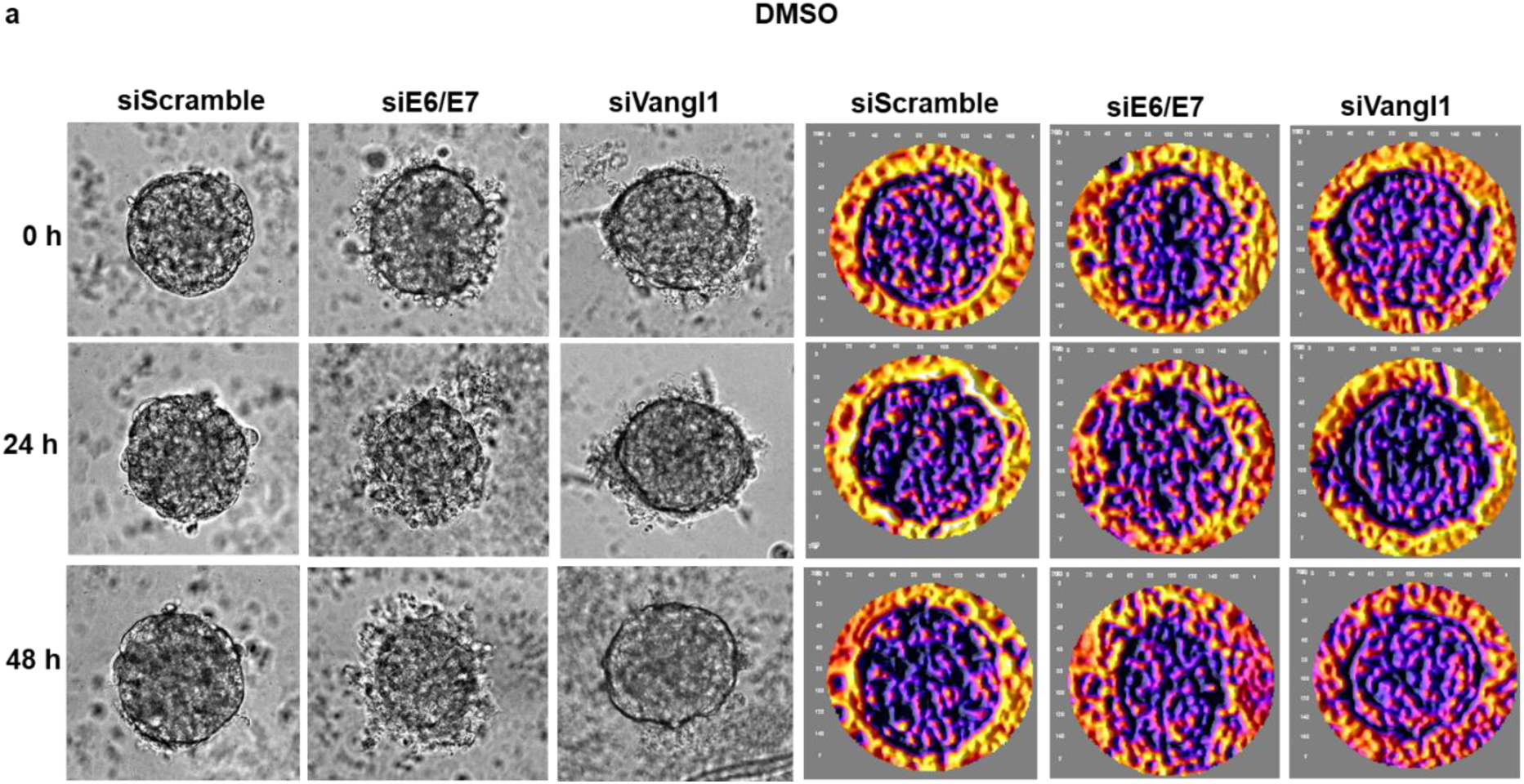
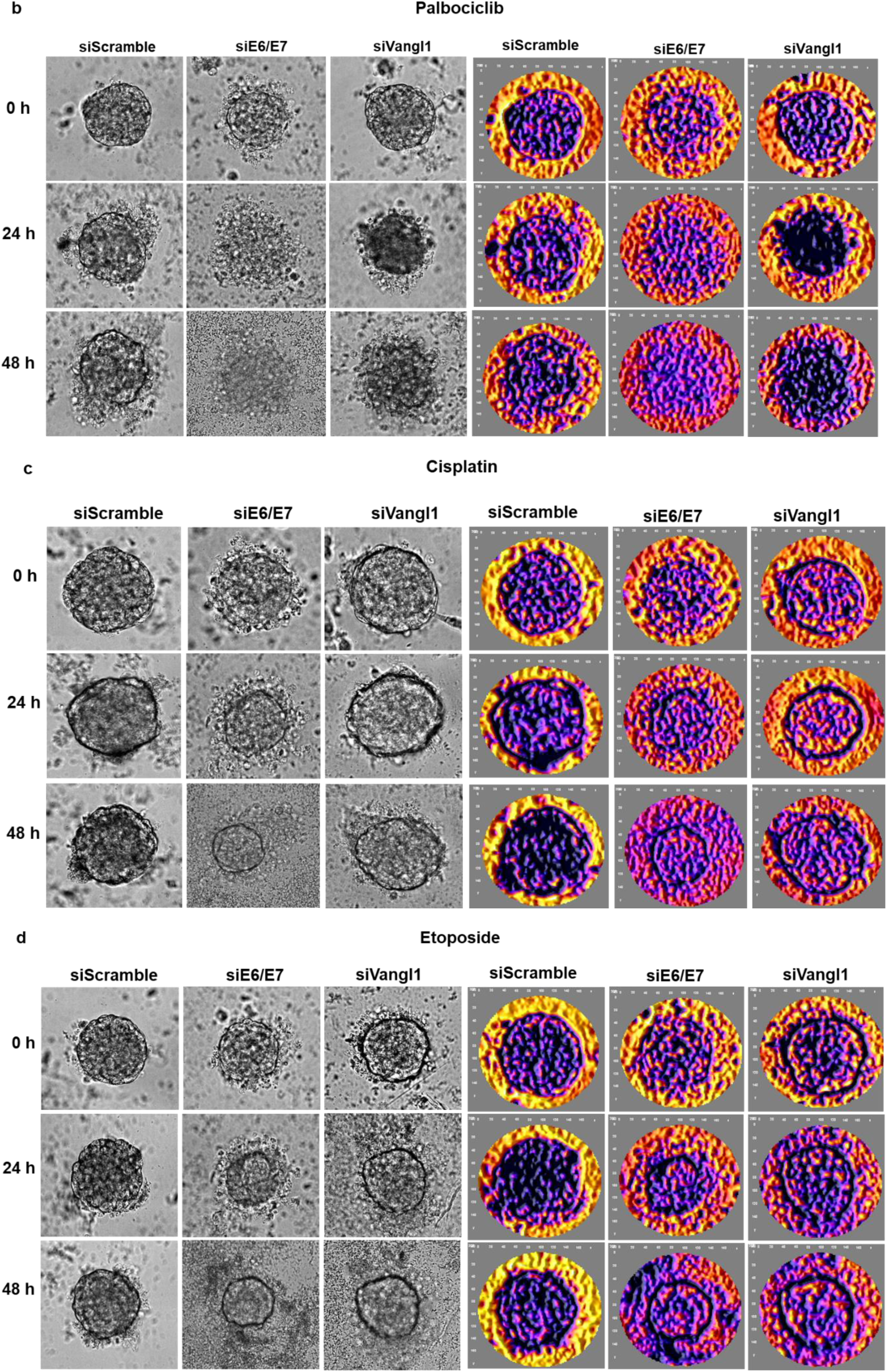
Loss of Vangl1 enhances the chemosensitivity of cervical cancer spheroids. (a) Brightfield images of CaSki spheroids following siRNA transfection and 48 h treatment with DMSO. Control spheroids remain compact and cohesive, whereas E6/E7- and Vangl1-depleted spheroids exhibit architectural changes; (b) Palbociclib treatment induces mild disintegration in siScramble spheroids, but causes pronounced disintegration in E6/E7- and Vangl1-depleted spheroids; (c–d) Cisplatin and Etoposide treatments show moderate disruption in siScramble spheroids, whereas E6/E7- and Vangl1-depleted spheroids undergo rapid dissociation and display debris-rich halos. The 3D surface plots of the spheroids are presented on the right panels. Brightfield images were acquired at 20x magnification. Micrographs and 3D surface plots were prepared using ImageJ. The brightness of the panels was uniformly enhanced post-image processing.

These findings show that depletion of E6/E7 or Vangl1 primes spheroids of transformed cervical cancer cells for treatment-induced disintegration. Taken together, we show that Vangl1 depletion enhances the sensitivity of CaSki spheroids to known chemotherapeutics used to treat cervical cancer.

## Discussion

High-risk HPV E7 proteins classically target the pRB tumour suppressor, but continuing research shows that E7 also influences cellular pathways beyond cell cycle control. In this study, we identify Vangl1, a core component of the Wnt/planar cell polarity (PCP) pathway (*72*), as a novel, phosphorylation-dependent interactor of HPV-16 E7. Overexpression or loss of Vangl1 has been reported to promote malignancy in breast cancer, glioblastoma multiforme, and lung adenocarcinoma (*66, 68, 73, 74*). Thus, the fate and role of Vangl1 in cancers is complex and context-dependent. This study therefore provides new insights into HPV-mediated cervical cancer by linking the E7 oncoprotein to a signalling axis that controls polarity and epithelial organization.

Our findings demonstrate that CKII phosphorylation of the HPV-16 E7 CR2 region is essential for Vangl1 binding. We found that Vangl1 binds strongly to phosphorylated HPV-16 E7, weakly to phosphorylated HPV-18 E7, and shows no detectable binding to the low-risk HPV E7 proteins examined. This specificity mirrors the oncogenic potential of these HPV subtypes (*75*) and suggests that recruitment of Vangl1 by HPV-16 may contribute to its pathogenicity. Additionally, the inability of phosphomimic peptides to recapitulate Vangl1-E7 binding further indicates that the interaction depends on actual structural features conferred by true phosphorylation rather than charge alone. CKII-mediated phosphorylation HPV-16 E7 at serine residues 31 and 32 (or 33 in HPV-18 E7) influences its interaction with known substrates like pRb, amplifying its degradation and consequently, cell transformability (*49*). Therefore, these observations establish a crucial biochemical requirement for Vangl1 recruitment and reiterate the importance of CKII phosphorylation on E7’s interactome.

Furthermore, our findings demonstrate that HPV-16 E7 specifically binds Vangl1 but not Vangl2. This confirms that Vangl1 can be regulated differently from Vangl2, despite their close structural similarities (*76*). A key insight from this study is that E7 not only binds Vangl1, but also modulates its abundance and turnover in HPV-transformed cervical cancer cells. Interestingly, this oncoprotein-specific modulation of Vangl1 levels is not fully explained by inhibition of canonical degradation pathways. Rather, cycloheximide chase assays showed a distinct biphasic post-translational decay of different Vangl1 forms, whereby unphosphorylated Vangl1 is rapidly degraded while there is an aberrant stabilization of the phosphorylated form. In addition, this prolonged retention of Vangl1 led to stabilization of E7. This bidirectional stabilization suggests that E7 contributes to the persistence of Vangl1 to, potentially, establish an oncogenic signalling. Studies have shown that high-risk E7 oncoprotein modulates several host factors for its own (e.g. pRB, USP7, USP11), or its the substrate’s stabilization (e.g. APOBEC3As) (*77–80*). These regulations occurring through direct interaction, through inhibiting degradation, or through a recently-described chaperone activity (with pRB), contribute towards establishing or sustaining malignancy in a cervical cancer context.

Mechanistically, our study indicates that E7 alters Vangl1 trafficking through the clathrin adaptor subunit AP1M1. In normal cells, AP1M1 is responsible for the endocytic sorting of Vangl1, facilitating its export from the TGN to the cell cortex, where it assembles into PCP complexes and interacts with cytoskeletal regulators (*60*). Therefore, sequestering AP1M1 can impair Vangl1 localization and function. We found that AP1M1 binds phosphorylated HPV-16 E7, and the effects of its depletion resemble those of E6/E7 knockdown, by increasing Vangl1 abundance and reducing E7 levels. Furthermore, E7 and AP1M1 jointly influence Vangl1 localization, disrupting its distribution to the cytoskeleton. Since Vangl1’s function as a PCP scaffold depends on its precise spatial distribution, E7-driven mislocalization disrupts polarity cues and contributes to the altered epithelial architecture characteristic of HPV-transformed cells. In addition, these observations further clarify the non-canonical Vangl1 rescue upon E7 depletion, as well as the prolonged half-life of Vangl1, as possible consequences of mislocalization. This is because altered localization can sequester proteins away from normal turnover and affect overall protein abundance (*56, 57*). This study also supports previous reports that CKII-phosphorylated E7 contributes to the mislocalization of PDZ-containing polarity proteins that are considered to be primary targets of the E6 oncoprotein (*81, 82*).

The functional consequences of this regulatory network are seen in CaSki spheroid models, where Vangl1 depletion phenocopies E6/E7 loss by inducing spheroid disintegration and loss of structural cohesion. These parallel phenotypes indicate that Vangl1 is not merely a passive target of E7, but an active effector required for maintaining epithelial integrity in HPV-positive cells. Beyond its role in maintaining spheroid architecture, Vangl1 is essential for the invasive capacity of transformed cervical cancer cells. Its loss produces distinct, quantifiable compromises in both architecture and invasive capacity, with the most pronounced defects emerging under conditions that normally support organized collective invasion. Indeed, Vangl1 knockdown markedly impaired invasion in both 2D and 3D models, underlining its requirement for efficient matrix engagement and directional movement. Of particular significance, Vangl1 depletion abolished the hallmark collective invasion phenotype exhibited by wild-type CaSki spheroids.

Although metastasis has traditionally been viewed as a predominantly single-cell process driven by loss of adhesion, accumulating evidence shows that tumour cells can migrate and disseminate as cohesive clusters that retain cell–cell contacts. Such collective invasion is directed by leader cells that extend into the surrounding matrix while remaining mechanically coupled to follower cells, generating contiguous invasive strands (*69*). This behaviour closely mirrors the invasive phenotype of wild-type CaSki spheroids in our study. Consistent with establishing the front-rear polarity cues required for this process(*71, 83*), we observed striking enrichment of Vangl1 at the leading edge of invasive protrusions, positioning it at the site where directional and collective coordination are established. These findings reinforce the idea that Vangl1 is a critical determinant of collective motility and a central effector through which E7 reshapes polarity and stabilizes oncogenic architecture in transformed cervical contexts.

In addition to its role in sustaining epithelial architecture and collective invasion, Vangl1 also proved essential for the ability of cervical cancer spheroids to withstand therapeutic stress. In response to treatment with all drugs (Etoposide, Cisplatin, and Palbociclib), the most severe structural deterioration occurred in spheroids lacking E6/E7, reinforcing the requirement for their continuous expression in sustaining the viability of cervical cancer cells. However, the comparable morphological effects seen upon Vangl1 depletion also indicate its contribution to maintaining viability under genotoxic and cytostatic stress; reinforcing the notion that Vangl1 functions as a downstream effector through which E7 maintains the structural resilience of transformed epithelial architecture. Moreover, Vangl1 depletion enhanced the cells’ chemosensitivity to drugs with distinct mechanisms of action, suggesting that Vangl1 contributes broadly to the cytostatic and genotoxic stress tolerance of cervical cancer cells. Our study therefore positions the Wnt/PCP pathway as a promising therapeutic axis for HPV-associated cervical cancer by identifying Vangl1 PCP signalling as a viable druggable target.

This study uncovers a novel role for HPV-16 E7 in modulating PCP signalling and identifies Vangl1 as a substrate in HPV-driven epithelial disorganization. Further research will determine how Vangl1 mislocalization influences downstream PCP effectors, the role of endocytic machineries in this, and how these changes contribute to cervical transformation in the HPV context. A deeper understanding of how HPV manipulates polarity signalling may reveal further new therapeutic targets.

## Materials and Methods

### Cell culture and reagents

HEK-293 cells, normal human keratinocytes, HaCaT cells, HPV-16/18–positive cervical cancer cell lines (CaSki and HeLa), and the HPV-negative cervical cancer line C-33A were obtained from ATCC. Cells were maintained in DMEM (Gibco, 31885-023) supplemented with 10% FBS (Gibco, 10270-106), penicillin–streptomycin (100 U mL^⁻¹^), and glutamine (300 μg mL^⁻¹^; Gibco, 10378-016). Cultures were grown at 37°C in a humidified incubator with 10% CO_₂_.

### Antibodies, inhibitors, antibiotics and chemotherapeutics

Primary antibodies included anti-Vangl1 (Merck, AMAB90600), anti-RB1/pRB (BD Pharmingen, 554136), anti-FLAG (Merck, F7425), anti-HA (Abcam, ab9110), anti-HPV-16 E7 (SCBT, sc65711), anti-phospho-HPV-16 E7 was generated by Eurogentec as previously described (ref), anti-HPV-18 E7 (SCBT, sc-365035), anti-AP1M1 (Merck, SAB1301057), anti-GAPDH (SCBT, sc-32233), anti-β-galactosidase (Promega, Z378B), anti-EGFR (SCBT, sc-373746), anti-Vimentin (SCBT, sc-6260), anti-Vinculin (CST, 4650T), HRP-conjugated anti-mouse (115-035-071) and anti-rabbit (111-035-046) secondary antibodies were from Jackson ImmunoResearch.

MG132 (Sigma-Aldrich, C2211) was used for proteasome inhibition, cycloheximide (Sigma-Aldrich, C4859) for protein synthesis inhibition, and chloroquine phosphate (Sigma-Aldrich, PHR1258) for lysosomal inhibition. CX-4945 (Silmitasertib) was used for inhibiting casein kinase II phosphorylation. Ampicillin (Merck, A5354) was used in bacterial cultures for selection.

Etoposide (Merck, E1383), Cisplatin (Merck, C2210000) and Palbociclib (CST, 47284) were used as chemotherapeutics.

### Plasmid constructs and transfections

pGEX-2T constructs encoding HPV-5 E7, HPV-11 E7, HPV-16 E7, and HPV-16 E7 C-terminal fragments were generated as described (*54*). Wild-type pGEX-18E7 and FLAG/HA-tagged pCMV16 E7 plasmids (*55*) were gifts from Karl Münger. E7 fragments were subcloned into pCMV GFP plasmid, a kind gift from Connie Cepko (Addgene,11153). The CKII α/β expression vector (pCDF CKII duet) was provided by Yong Xiong and John Guatelli. FLAG-Vangl1 was provided by Pablo Rodriguez-Viciana (*56*) and Vangl2 ORF was cloned into pcDNA3.1 (Genscript). All constructs were sequence-verified.

HEK-293 cells were transfected using the calcium phosphate precipitation method (*57*).

### Peptide synthesis

Peptides were synthesized at the ICGEB Biotechnology Development Unit. Scrambled peptides contained 10 random amino acids. Non-phosphorylated peptides corresponded to HPV-16 E7 CR2 (residues 15–39; wild-type or N29S). Phosphorylated peptides contained serine phosphorylation at S29, S31, and S32.

### Peptide pulldown assay

HaCaT cells (80–90% confluent) were lysed in RIPA buffer containing 50 mM Tris-Cl pH 7.4, 150 mM NaCl, 1% NP-40, 0.5% sodium deoxycholate, 0.1% SDS, and protease inhibitors. Lysates were clarified by centrifugation (16,000 rpm, 10 min, 4°C). Biotinylated peptides (500 μg) were bound to streptavidin magnetic beads (GE Healthcare) for 1 h at 4°C. Pre-cleared lysates were incubated with peptide-bound beads for 2 h at 4°C. After extensive washing, 5% of the beads were analyzed by Western blotting; the remainder was digested with trypsin and analyzed by mass spectrometry.

### Mass spectrometry

Protein complexes were analyzed as described previously (*58*). Samples were loaded onto Picofrit nanobore columns packed with 1.8 μm Zorbax XDB C18 resin and analyzed on an LTQ ion trap mass spectrometer (Thermo Electron). One full MS scan (400–1700 m/z) was followed by eight data-dependent MS/MS scans. RAW files were converted to mzXML (READW v1.6) and searched against Ensembl human and NCBInr viral databases using the Global Proteome Machine with X!Tandem.

### GST-fusion protein purification

pGEX-2T constructs (GST empty, GST-5E7, GST-11E7, GST-16E7, GST-16E7 C-T, GST-18E7) were transformed into *E. coli* BL21(DE3) pLysS. For phosphorylated GST-E7 proteins, constructs were co-transformed with pCDF-CKII. Cultures were induced with 200 μM IPTG at 25°C for 16–20 h. Bacteria were lysed in PBS with 1% Triton X-100, sonicated, clarified, and incubated with glutathione agarose beads (Sigma, G4510). Beads were washed and stored in PBS with 1% Triton X-100 and 30% glycerol at −20°C.

### GST pull-down assay

HaCaT cells were lysed in RIPA buffer on ice for 10 min and clarified by centrifugation. Lysates were incubated with GST-fusion proteins for 2 h at 4°C. Beads were washed with E1A buffer (50 mM HEPES pH 7.0, 250 mM NaCl, 0.1% NP-40), eluted in 2× SDS loading buffer, and analyzed by SDS-PAGE and Western blotting.

### Co-immunoprecipitation

HEK293 cells were transfected as indicated and lysed in RIPA buffer after 18 h. Clarified lysates were incubated with EZview Red Anti-HA Affinity Gel (Sigma, E6779) for 4 h at 4°C. Beads were washed, eluted in 2× SDS loading buffer, and analyzed by SDS-PAGE and Western blotting.

### siRNA transfection

CaSki, HeLa, C-33A and HaCaT cells were transfected with siRNAs targeting Vangl1 (Dharmacon SMARTpool), AP1M1 (Dharmacon SMARTpool), HPV-16 E6/E7 (Eurofins), or Scramble control (Dharmacon siSTABLE) as previously described (*52*). Cells were harvested after 72 h for Western blotting or downstream assays.

For spheroids, siRNA mixes were prepared in serum-free DMEM, but spheroids were maintained in DMEM with FBS and PSG. Partial media replacement was performed after 24 h to preserve spheroid integrity.

### Western blotting

Samples were denatured at 95°C for 10 min, separated by SDS-PAGE, and transferred to 0.22-µm nitrocellulose membranes. Membranes were blocked in 5% milk/TBS-T for 1 h, incubated with primary and secondary antibodies, and developed using ECL (GE Healthcare, RPN2106).

### Proteasomal and lysosomal inhibition

Cells transfected with siRNAs were treated with 20 µM MG132 or chloroquine for 6 h (endogenous proteins) after 72 h. Then, cells were harvested for Western blot analysis.

### Cycloheximide chase

CaSki cells transfected with siRNA for 72 h were treated with 50 µg/mL cycloheximide and harvested at defined time points (0–10 h). HEK293 cells overexpressing Vangl1, HPV-16 E7, or both were treated similarly, after 24 h of transfection. Protein stability was assessed by Western blotting.

### Two-dimensional Matrigel invasion assay

Matrigel invasion chambers (Corning BioCoat) were rehydrated with serum-free DMEM for 2 h. siRNA-transfected CaSki cells (5 × 10⁴) were seeded into the upper chamber in serum-free DMEM; the lower chamber contained DMEM with 10% FBS. After 20 h, invading cells were fixed, permeabilized, stained with 0.05% Crystal Violet, and imaged at 10× magnification. At least five fields were analyzed per sample.

### Generation of 3D CaSki spheroids

Spheroids were generated by seeding 5,000 CaSki cells in 100 µL per well in ULA 96-well round-bottom plates. Plates were incubated with gentle agitation at 37°C and 10% CO₂. Compact spheroids formed within 48 h, with diameters of 160–170 µm. Spheroids were further incubated for 48 h before being used for assays.

### Collagen invasion assay

Coverslips were coated with 1 mg/mL poly-L-lysine. Collagen I was neutralized to pH 7.0, and Matrigel–collagen mixtures (1:0, 1:1, 1:3) were prepared. Twenty microliters of matrix were placed on coverslips. Spheroids were transferred into the matrix and polymerized for 30 min at 37°C. Wells were filled with DMEM + 5% FBS. Invasion was monitored for 7 days and imaged at 20× magnification using Nikon Eclipse Ti2 inverted microscope equipped with the NIS-Elements AR 5.02.

### Immunofluorescence

Spheroids embedded in collagen were fixed with 4% paraformaldehyde for 20 min, permeabilized overnight at 4°C in 0.5% Triton X-100, and blocked in PBS containing 5% horse serum, 0.1% Triton X-100 for 2 h. Samples were incubated with anti-Vangl1 overnight at 4°C, washed, incubated with Alexa Fluor® 594 secondary antibody for 2 h. Cell nuclei were stained with DAPI and imaged using a Zeiss LSM 880 Airyscan microscope at 63× magnification. Image analysis was performed using ImageJ v1.54f.

HEK293 cells expressing GFP-16 E7 and FLAG-Vangl1 on poly-L-lysine coated coverslips were fixed with 4% paraformaldehyde for 10 min, permeabilized for 10 min in 0.1% Triton X-100, and blocked in PBS containing 1% bovine serum albumin for 30 min. Samples were incubated with anti-FLAG overnight at 4°C, washed, incubated with Alexa Fluor® 594 secondary antibody for 1 h. Nuclei staining and imaging were done as mentioned above.

### Chemotherapy treatment of spheroids

CaSki spheroids pre-transfected with siRNAs for 48 h were treated with DMSO (as control), cisplatin, etoposide or palbociclib. Spheroids were imaged at 24 h and 48 h post treatment using Nikon Eclipse Ti2 inverted microscope equipped with the NIS-Elements AR 5.02.

### Statistical analysis

Statistical analyses were performed using GraphPad Prism 10. Comparisons between control and siRNA-treated cells were made using Student’s *t*-test. Significance was defined as *p* < 0.05. Experiments were performed at least three times, and data are presented as mean ± standard deviation. Significance is indicated as: *p* < 0.05 \**, p < 0.005 ***, p < 0.001 \*\*\*\**, and non-significant (ns) for *p* > 0.05.

Protein quantification was performed using ImageJ v2.16/1.54p, with relative levels calculated as the ratio of target band intensity to loading-control intensity.

## Acknowledgments

We thank Miranda Thomas for her valuable comments on the manuscript. IDG is a recipient of the WeSTAR fellowship of the Italian Ministry of Foreign Affairs and International Cooperation and the Postdoctoral fellowship of Standing Committee for Scientific and Technological Cooperation (COMSTECH). GDS isa recipient of the Fondazione AIRC IG grant 30570, the Fondazione AIRC Special Programme Molecular Clinical Oncology “5 per mille” 22759 and the Worldwide Cancer Research (grant 24-0361). RB is supported by an AIRC post-doctoral fellowship for Italy.

## Author contributions

Conceptualization: IDG, OB, LB

Methodology: IDG, OB, MPM, RB, GDS, LB

Investigation: IDG, OB, MPM, RB, LB

Visualization: IDG, OB

Supervision: LB

Writing—original draft: IDG, LB

Writing—review & editing: IDG, OB, MPM, RB, GDS, LB

## Data and materials availability

All data supporting the findings of this study are included within the article or are available from the authors upon reasonable request.

## References

1. World Health Organization, Cervical cancer, https://www.who.int/news-room/fact-sheets/detail/cervical-cancer#:~:text=HPV%20vaccination%20and%20other%20prevention,HPV%20infection%20and%20cervical%20cancer. (2024).

2. R. U. Hakim, T. Amin, S. M. B. Ul Islam, Advances and Challenges in Cervical Cancer: From Molecular Mechanisms and Global Epidemiology to Innovative Therapies and Prevention Strategies. Cancer Control 32 (2025).

3. E. M. Burd, Human Papillomavirus and Cervical Cancer. Clin. Microbiol. Rev. 16, 1–17 (2003).

4. S. F. Jabbar, L. Abrams, A. Glick, P. F. Lambert, Persistence of high-grade cervical dysplasia and cervical cancer requires the continuous expression of the human papillomavirus type 16 E7 oncogene. Cancer Res. 69 (2009).

5. R. J. Kurman, M. H. Schiffman, W. D. Lancaster, R. Reid, A. B. Jenson, G. F. Temple, A. T. Lorincz, Analysis of individual cervical human papillomavirus types in neolasia: A possible role for type 18 in rapid progression. Am. J. Obstet. Gynecol. 159 (1988).

6. M. E. Scheurer, G. Tortolero-Luna, M. Guillaud, M. Follen, Z. Chen, L. M. Dillon, K. Adler-Storthz, Correlation of human papillomavirus type 16 and human papillomavirus type 18 E7 messenger RNA levels with degree of cervical dysplasia. Cancer Epidemiology Biomarkers and Prevention 14 (2005).

7. S. F. Jabbar, S. Park, J. Schweizer, M. Berard-Bergery, H. C. Pitot, D. Lee, P. F. Lambert, Cervical cancers require the continuous expression of the human papillomavirus type 16 E7 oncoprotein even in the presence of the viral E6 oncoprotein. Cancer Res. 72 (2012).

8. C. K. Choo, E. A. Rorke, R. L. Eckert, Differentiation-independent Constitutive Expression of the Human Papillomavirus Type 16 E6 and E7 Oncogenes in the CaSki Cervical Tumour Cell Line. Journal of General Virology 75, 1139–1147 (1994).

9. L. Zhou, Q. Qiu, Q. Zhou, J. Li, M. Yu, K. Li, L. Xu, X. Ke, H. Xu, B. Lu, H. Wang, W. Lu, P. Liu, Y. Lu, Long-read sequencing unveils high-resolution HPV integration and its oncogenic progression in cervical cancer. Nat. Commun. 13 (2022).

10. L. Banks, P. Spence, E. Androphy, N. Hubbert, G. Matlashewski, A. Murray, L. Crawford, Identification of human papillomavirus type 18 E6 polypeptide in cells derived from human cervical carcinomas. Journal of General Virology 68 (1987).

11. E. Schwarz, U. K. Freese, L. Gissmann, W. Mayer, B. Roggenbuck, A. Stremlau, H. Zur Hausen, Structure and transcription of human papillomavirus sequences in cervical carcinoma cells. Nature 314 (1985).

12. C. Y. Zhang, W. Bao, L. H. Wang, Downregulation of p16ink4a inhibits cell proliferation and induces G1 cell cycle arrest in cervical cancer cells. Int. J. Mol. Med. 33 (2014).

13. J. Wang, A. Sampath, P. Raychaudhuri, S. Bagchi, Both Rb and E7 are regulated by the ubiquitin proteasome pathway in HPV-containing cervical tumour cells. Oncogene 20 (2001).

14. E. A. Harrington, J. L. Bruce, E. Harlow, N. Dyson, pRB plays an essential role in cell cycle arrest induced by DNA damage. Proc. Natl. Acad. Sci. U. S. A. 95 (1998).

15. M. S. Barbosa, C. Edmonds, C. Fisher, J. T. Schiller, D. R. Lowy, K. H. Vousden, The region of the HPV E7 oncoprotein homologous to adenovirus E1a and Sv40 large T antigen contains separate domains for Rb binding and casein kinase II phosphorylation. EMBO J. 9 (1990).

16. O. Basukala, S. Mittal, P. Massimi, M. Bestagno, L. Banks, The HPV-18 E7 CKII phospho acceptor site is required for maintaining the transformed phenotype of cervical tumour-derived cells. PLoS Pathog. 15 (2019).

17. M. P. Dizanzo, M. Bugnon Valdano, O. Basukala, L. Banks, D. Gardiol, Novel effect of the high risk-HPV E7 CKII phospho-acceptor site on polarity protein expression. BMC Cancer 22 (2022).

18. A. Javorsky, P. O. Humbert, M. Kvansakul, Viral manipulation of cell polarity signalling. [Preprint] (2023). 10.1016/j.bbamcr.2023.119536.

19. L. Banks, D. Pim, M. Thomas, Human tumour viruses and the deregulation of cell polarity in cancer. [Preprint] (2012). 10.1038/nrc3400.

20. M. P. Dizanzo, F. Marziali, C. Brunet Avalos, M. Bugnon Valdano, S. Leiva, A. L. Cavatorta, D. Gardiol, HPV E6 and E7 oncoproteins cooperatively alter the expression of Disc Large 1 polarity protein in epithelial cells. BMC Cancer 20 (2020).

21. Y. Yang, M. Mlodzik, Wnt-Frizzled/Planar Cell Polarity Signaling: Cellular Orientation by Facing the Wind (Wnt). Annu. Rev. Cell Dev. Biol. 31 (2015).

22. W. Yang, L. Garrett, D. Feng, G. Elliott, X. Liu, N. Wang, Y. M. Wong, N. T. Choi, Y. Yang, B. Gao, Wnt-induced Vangl2 phosphorylation is dose-dependently required for planar cell polarity in mammalian development. Cell Res. 27 (2017).

23. C. A. Dreyer, K. VanderVorst, K. L. Carraway, Vangl as a Master Scaffold for Wnt/Planar Cell Polarity Signaling in Development and Disease. [Preprint] (2022). 10.3389/fcell.2022.887100.

24. K. VanderVorst, C. A. Dreyer, J. Hatakeyama, G. R. R. Bell, J. A. Learn, A. L. Berg, M. Hernandez, H. Lee, S. R. Collins, K. L. Carraway, Vangl-dependent Wnt/planar cell polarity signaling mediates collective breast carcinoma motility and distant metastasis. Breast Cancer Research 25 (2023).

25. L. Zhao, L. Wang, C. Zhang, Z. Liu, Y. Piao, J. Yan, R. Xiang, Y. Yao, Y. Shi, E6-induced selective translation of WNT4 and JIP2 promotes the progression of cervical cancer via a noncanonical WNT signaling pathway. Signal Transduct. Target. Ther. 4 (2019).

26. R. Zhang, H. Lu, Y. Y. Lyu, X. M. Yang, L. Y. Zhu, G. D. Yang, P. C. Jiang, Y. Re, W. W. Song, J. H. Wang, C. C. Zhang, F. Gu, T. J. Luo, Z. Y. Wu, C. J. Xu, E6/E7-P53-POU2F1-CTHRC1 axis promotes cervical cancer metastasis and activates Wnt/PCP pathway. Sci. Rep. 7 (2017).

27. C. A. Dreyer, K. VanderVorst, D. Natwick, G. Bell, P. Sood, M. Hernandez, J. M. Angelastro, S. R. Collins, K. L. Carraway, A complex of Wnt/planar cell polarity signaling components Vangl1 and Fzd7 drives glioblastoma multiforme malignant properties. Cancer Lett. 567 (2023).

28. J. Hatakeyama, J. H. Wald, I. Printsev, H. Y. H. Ho, K. L. Carraway, Vangl1 and Vangl2: Planar cell polarity components with a developing role in cancer. Endocr. Relat. Cancer 21 (2014).

29. A. Zine El Abidine, V. Tomaić, R. Bel Haj Rhouma, P. Massimi, I. Guizani, S. Boubaker, E. Ennaifer, L. Banks, A naturally occurring variant of HPV-16 E7 exerts increased transforming activity through acquisition of an additional phospho-acceptor site. Virology 500 (2017).

30. L. B. Chemes, I. E. Sánchez, C. Smal, G. De Prat-Gay, Targeting mechanism of the retinoblastoma tumour suppressor by a prototypical viral oncoprotein: Structural modularity, intrinsic disorder and phosphorylation of human papillomavirus E7. FEBS Journal 277 (2010).

31. D. Feng, J. Wang, W. Yang, J. Li, X. Lin, F. Zha, X. Wang, L. Ma, N. T. Choi, Y. Mii, S. Takada, M. S. Y. Huen, Y. Guo, L. Zhang, B. Gao, Regulation of Wnt/PCP signaling through p97/VCP-KBTBD7-mediated Vangl ubiquitination and endoplasmic reticulum-associated degradation. Sci. Adv. 7 (2021).

32. R. S. Hegde, E. Zavodszky, Recognition and degradation of mislocalized proteins in health and disease. Cold Spring Harb. Perspect. Biol. 11 (2019).

33. S. Mallik, S. Kundu, Topology and Oligomerization of Mono- and Oligomeric Proteins Regulate Their Half-Lives in the Cell. Structure 26 (2018).

34. M. Yasumura, A. Hagiwara, Y. Hida, T. Ohtsuka, Planar cell polarity protein Vangl2 and its interacting protein Ap2m1 regulate dendritic branching in cortical neurons. Genes to Cells 26 (2021).

35. P. Navarro Negredo, J. R. Edgar, A. G. Wrobel, N. R. Zaccai, R. Antrobus, D. J. Owen, M. S. Robinson, Contribution of the clathrin adaptor AP-1 subunit µ1 to acidic cluster protein sorting. Journal of Cell Biology 216, 2927–2943 (2017).

36. J. M. Carvajal-Gonzalez, S. Balmer, M. Mendoza, A. Dussert, G. Collu, A. C. Roman, U. Weber, B. Ciruna, M. Mlodzik, The clathrin adaptor AP-1 complex and Arf1 regulate planar cell polarity in vivo. Nat. Commun. 6 (2015).

37. K. Mizuguchi, H. Aoki, M. Aoyama, Y. Kawaguchi, Y. Waguri-Nagaya, N. Ohte, K. Asai, Three-dimensional spheroid culture induces apical-basal polarity and the original characteristics of immortalized human renal proximal tubule epithelial cells. Exp. Cell Res. 404 (2021).

38. M. E. Feigin, S. D. Akshinthala, K. Araki, A. Z. Rosenberg, L. B. Muthuswamy, B. Martin, B. D. Lehmann, H. K. Berman, J. A. Pietenpol, R. D. Cardiff, S. K. Muthuswamy, Mislocalization of the cell polarity protein scribble promotes mammary tumourigenesis and is associated with basal breast cancer. Cancer Res. 74 (2014).

39. F. Peglion, S. Etienne-Manneville, Cell polarity changes in cancer initiation and progression. [Preprint] (2024). 10.1083/jcb.202308069.

40. C. Laumonnerie, D. J. Solecki, Regulation of Polarity Protein Levels in the Developing Central Nervous System. [Preprint] (2018). 10.1016/j.jmb.2018.05.036.

41. S. V. Paramore, C. Trenado-Yuste, R. Sharan, C. M. Nelson, D. Devenport, Vangl-dependent mesenchymal thinning shapes the distal lung during murine sacculation. Dev. Cell 59, 1302–1316.e5 (2024).

42. J. N. Anastas, T. L. Biechele, M. Robitaille, J. Muster, K. H. Allison, S. Angers, R. T. Moon, A protein complex of SCRIB, NOS1AP and VANGL1 regulates cell polarity and migration, and is associated with breast cancer progression. Oncogene 31 (2012).

43. J. H. Wald, J. Hatakeyama, I. Printsev, A. Cuevas, W. H. D. Fry, M. J. Saldana, K. Vandervorst, A. Rowson-Hodel, J. M. Angelastro, C. Sweeney, K. L. Carraway, Suppression of planar cell polarity signaling and migration in glioblastoma by Nrdp1-mediated Dvl polyubiquitination. Oncogene 36 (2017).

44. A. A. Khalil, P. Friedl, Determinants of leader cells in collective cell migration. Integrative Biology 2, 568 (2010).

45. K. V. Nguyen-Ngoc, K. J. Cheung, A. Brenot, E. R. Shamir, R. S. Gray, W. C. Hines, P. Yaswen, Z. Werb, A. J. Ewald, ECM microenvironment regulates collective migration and local dissemination in normal and malignant mammary epithelium. Proc. Natl. Acad. Sci. U. S. A. 109 (2012).

46. A. Gandalovičová, T. Vomastek, D. Rosel, J. Brábek, Cell polarity signaling in the plasticity of cancer cell invasiveness. Oncotarget 7 (2016).

47. C. cheng Hao, C. yang Xu, X. yu Zhao, J. ning Luo, G. Wang, L. hong Zhao, X. Ge, X. feng Ge, Up-regulation of VANGL1 by IGF2BPs and miR-29b-3p attenuates the detrimental effect of irradiation on lung adenocarcinoma. Journal of Experimental and Clinical Cancer Research 39 (2020).

48. Y. Y. J. Z. Y. Z. S. Z. J. C. W. M. Xin Feng, Core planar cell polarity genes VANGL1 and VANGL2 in predisposition to congenital vertebral malformations. PNAS (2024).

49. C. H. Lin, H. S. Chang, W. C. Y. Yu, USP11 stabilizes HPV-16E7 and further modulates the E7 biological activity. Journal of Biological Chemistry 283, 15681–15688 (2008).

50. J. A. Westrich, C. J. Warren, M. J. Klausner, K. Guo, C.-W. Liu, M. L. Santiago, D. Pyeon, Human Papillomavirus 16 E7 Stabilizes APOBEC3A Protein by Inhibiting Cullin 2-Dependent Protein Degradation. J. Virol. 92 (2018).

51. C. Xia, C. Xiao, H. Y. Luk, P. K. S. Chan, S. S. Boon, The ubiquitin specific protease 7 stabilizes HPV16E7 to promote HPV-mediated carcinogenesis. Cellular and Molecular Life Sciences 80 (2023).

52. I. Gbala, N. Kavcic, L. Banks, The retinoblastoma protein contributes to maintaining the stability of HPV E7 in cervical cancer cells. J. Virol. 99 (2025).

53. J. shun Wu, J. Jiang, B. jun Chen, K. Wang, Y. ling Tang, X. hua Liang, Plasticity of cancer cell invasion: Patterns and mechanisms. [Preprint] (2021). 10.1016/j.tranon.2020.100899.

54. D. Pim, P. Massimi, S. M. Dilworth, L. Banks, Activation of the protein kinase B pathway by the HPV-16 E7 oncoprotein occurs through a mechanism involving interaction with PP2A. Oncogene 24 (2005).

55. S. L. Gonzalez, M. Stremlau, X. He, J. R. Basile, K. Münger, Degradation of the Retinoblastoma Tumour Suppressor by the Human Papillomavirus Type 16 E7 Oncoprotein Is Important for Functional Inactivation and Is Separable from Proteasomal Degradation of E7. J. Virol. 75 (2001).

56. L. C. Young, N. Hartig, M. Muñoz-Alegre, J. A. Oses-Prieto, S. Durdu, S. Bender, V. Vijayakumar, M. VietriRudan, C. Gewinner, S. Henderson, A. P. Jathoul, R. Ghatrora, M. F. Lythgoe, A. L. Burlingame, P. Rodriguez-Viciana, An MRAS, SHOC2, and SCRIB complex coordinates erk pathway activation with polarity and tumourigenic growth. Mol. Cell 52 (2013).

57. C. Chen, H. Okayama, High-efficiency transformation of mammalian cells by plasmid DNA. Mol. Cell. Biol. 7 (1987).

58. V. Tomaić, D. Gardiol, P. Massimi, M. Ozbun, M. Myers, L. Banks, Human and primate tumour viruses use PDZ binding as an evolutionarily conserved mechanism of targeting cell polarity regulators. Oncogene 28 (2009).

